# Dietary fiber selectively regulates intestinal persistence of probiotic bifidobacteria

**DOI:** 10.64898/2026.06.06.730603

**Authors:** Garam Choi, Shruti Chatterjee, Urmi Shah, Tae Hyung Won, Alexandros Skouris, Jaehyeon Kim, Waleed Mujib, Haipeng Sun, Marissa A. Fontaine, Adriana Messyasz, Alexander Lemenze, David K. Meyerholz, María G. Domínguez-Bello, Frank C. Schroeder, Douwe van Sinderen, Lok Yin Roy Wong, Nicholas J. Bessman

**Affiliations:** Center for Immunity and Inflammation, Rutgers New Jersey Medical School, Newark, NJ, USA; Department of Medicine, Rutgers New Jersey Medical School, Newark, NJ, USA; Center for Virus-Host Innate Immunity, Rutgers New Jersey Medical School, Newark, NJ, USA; Department of Microbiology, Biochemistry and Molecular Genetics, Rutgers New Jersey Medical School, Newark, NJ, USA; Department of Chemistry and Chemical Biology, Boyce Thompson Institute, Cornell University, Ithaca, NY, USA; Department of Biochemistry & Microbiology, Rutgers University, New Brunswick, NJ, USA; Molecular and Genomics Informatics Core Facility, Rutgers New Jersey Medical School, Newark, NJ, USA; Department of Pathology, Immunology and Laboratory Medicine, Rutgers New Jersey Medical School, Newark, NJ, USA; Department of Pathology, Roy J. and Lucille A. Carver College of Medicine, University of Iowa, Iowa City, IA, USA; APC Microbiome Ireland and School of Microbiology, University College Cork, Ireland; Department of Pharmacology, Sungkyunkwan University School of Medicine, Suwon, Republic of Korea

## Abstract

Bifidobacteria dominate the gut microbiota of breast-fed infants, and are strongly associated with human health, including immune regulation, colonization resistance, and protection against inflammation. However, bifidobacteria persist at high abundance after weaning in only a subset of individuals, and the factors that regulate intestinal persistence of bifidobacteria are poorly understood. Using gnotobiotic mouse models, we identified a common dietary fiber, raffinose, as a critical determinant of bifidobacterial persistence during microbial transitions associated with weaning. Bifidobacterial persistence depends on an intact raffinose utilization operon and is associated with disease resistance and restrained inflammation in adult mice. Specific dietary fiber recommendations commencing at weaning are a potential strategy to maintain bifidobacteria persistence beyond infancy, with potential long-term benefits for host resilience and reduced risk of inflammatory disease.

## Main

*Bifidobacterium* is the keystone genus in the breastfed infant gut^1,2^. As an early colonizer of the human gut, intestinal bifidobacteria uniquely promote healthy infant weight gain^3,4^, intestinal barrier function^5^, and resolution of inflammation^6^, prevent infection^7,8^, and enhance the efficacy of cancer immunotherapy^9,10^. Recent human studies further showed that bifidobacteria limit systemic inflammation^11^, promote vaccine efficacy^12^, and protect against influenza^13^. Despite its critical contribution to host health, bifidobacteria undergo a marked ecological transition during weaning when the infant microbiota diversifies and dietary substrates change. While some individuals maintain relatively high levels of bifidobacteria throughout childhood and adulthood, reaching up to 10-30% of the intestinal microbiota, others experience a dramatic decline or complete loss of these organisms^8,14–16^.

The intestinal abundance of commensal bifidobacteria can be influenced by multiple factors, including human milk oligosaccharides (HMOs)^17–20^, prebiotic dietary fibers^21–26^, and dietary iron supplementation^6,27^. However, while early-life bifidobacterial dominance is primarily driven by HMO-dependent priority effects during breastfeeding^11,28^, the factors that determine whether bifidobacteria persist or disappear after weaning remain poorly understood^29^. This question represents a major unresolved problem in human microbiome ecology, because long-term persistence of beneficial microbes may be important for sustaining their effects on host health. Current evidence from clinical studies in which HMOs or other prebiotics are administered demonstrates substantial inter-individual variability in bifidobacterial responses, and global analyses further reveal significant geographic variation in their persistence^8,16,19^. Together, these observations suggest that bifidobacterial persistence is shaped not only by early colonization history, but also by ongoing ecological interactions with the host diet and surrounding microbial community. Identifying factors that selectively support bifidobacterial persistence and abundance, and understanding their underlying mechanisms, could provide new strategies to improve human health outcomes.

To this end, we developed gnotobiotic mouse models that enable mechanistic investigation in a simplified microbial environment. While bifidobacteria colonize poorly in conventional mice^30–32^, this system uniquely allows controlled assessment of colonization dynamics. We selected *Bifidobacterium breve* as a model species in our study, since it frequently colonizes human infants^2,33^, and has been shown to confer protection in adult mouse models for immunotherapy response^9^, long-term pathogen colonization resistance^33^, and dextran sodium sulfate (DSS)-induced colitis^34^.

First, germ-free mice were mono-colonized with *B. breve* strain DSM 20213 (*Bb* hereafter) to examine whether host adaptive immunity or dietary iron directly modulates its intestinal persistence. However, neither the host adaptive immune system (Extended Data Fig. 1a), nor modified dietary iron levels (Extended Data Fig. 1b), affected *Bb* persistent colonization levels in mono-colonized mice. By contrast, when mice were switched from a grain-based standard lab diet (STD) to a chemically-defined AIN-93G diet containing non-fermentable cellulose fiber^35^, either with (AIN+IP) or without (AIN) supplemental inulin and pectin fibers, fecal *Bb* colony-forming units (CFUs) were reduced by at least 10-fold regardless of dietary iron levels (Fig. 1a and Extended Data Fig. 1b,c). Although some human studies suggest that inulin and/or pectin fiber can increase the relative abundance of bifidobacteria in some patients^23–26^, these fibers did not prevent the loss of *Bb* in this context (Extended Data Fig. 1b,c).

**Fig. 1.**
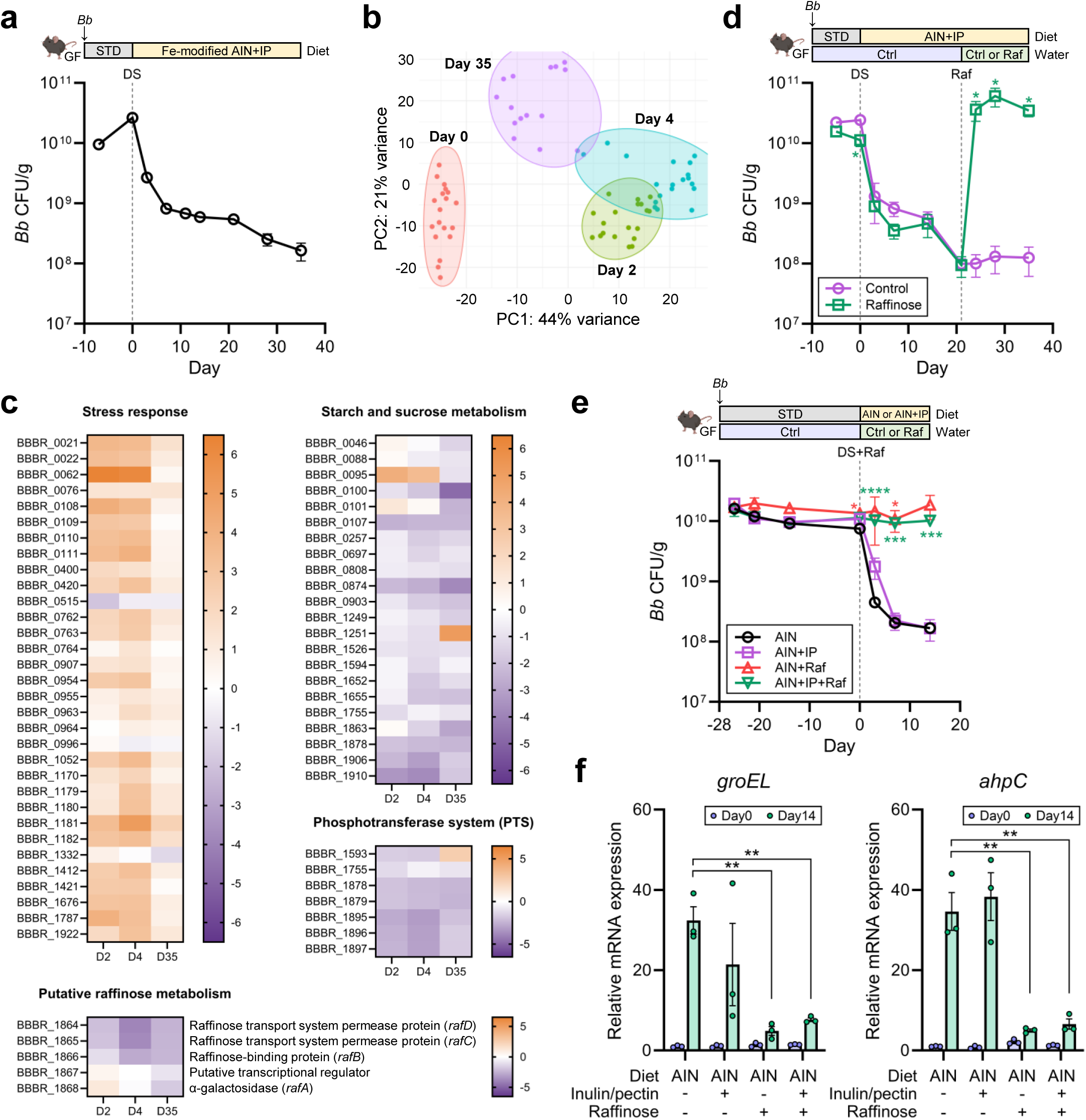
Intestinal colonization of *Bb* is impaired by switch to AIN-93G diets, and rescued by raffinose in mono-colonized mice. **a-c**, Germ-free mice were mono-colonized with *Bb*, followed by a switch from standard lab diet (STD) to Fe-modified AIN-93G diets supplemented with 1% inulin and 1% pectin (AIN+IP). Both low-Fe and high-Fe AIN+IP were used and pooled for analysis because they showed comparable effects on *Bb* colonization and transcriptomes. Fecal *Bb* CFUs were quantified over time by plating (**a**), and transcriptomic changes in *Bb* at day 0, 2, 4, and 35 after diet switch were analyzed by RNA-seq (**b,c**). Data were pooled from two independent experiments (*n* = 19). **b**, Principal component analysis (PCA) of *Bb* transcriptomes upon diet switch. Each point represents an individual sample and is colored by time points. **c**, Heatmap visualizing the relative expression of genes belonging to significantly altered pathways (as identified in Extended Data Fig. 2b). Gene expression levels at day 2, 4, and 35 were normalized to day 0, using a log_2_ scale. **d**, Germ-free mice were mono-colonized with *Bb*, followed by a switch from STD to AIN+IP. After 3 weeks on AIN+IP, mice were given either control drinking water or drinking water supplemented with 6.5% (w/v) raffinose. Fecal *Bb* CFUs were quantified over time by plating. Data represent *n* = 3 per group from one experiment. **e**,**f**, Germ-free mice were mono-colonized with *Bb* and maintained on STD for 4 weeks. Then, mice were switched to either AIN or AIN+IP with or without 1% (w/v) raffinose-supplemented water. Data represent *n* = 3 for AIN, *n* = 3 for AIN+IP, *n* = 2-3 for AIN+Raf, and *n* = 3-4 for AIN+IP+Raf. **e**, Fecal *Bb* CFUs were quantified over time by plating. Data are shown as mean ± SEM, and each group was compared to the AIN group by Student’s *t*-test (\**p* < 0.05; \*\*\**p* < 0.0005; \*\*\*\**p* < 0.0001). **f**, Expression levels of stress-response genes in *Bb* at day 0 and day 14 after a diet switch were analyzed by qPCR. Relative expression was normalized to the AIN without raffinose group at day 0 of the diet switch. Data in **a**,**d,f** are shown as mean ± SEM and were compared by Student’s *t*-test (\**p* < 0.05; \*\**p* < 0.005).

To better understand how the switch from STD to AIN+IP diet leads to decreased *Bb* persistence in mono-colonized mice, longitudinal fecal *Bb* RNA was isolated and analyzed by RNA-seq. We observed a clear and rapid transcriptome shift over time after the diet switch (Fig. 1b and Extended Data Fig. 2a). In particular, genes related to stress response^36,37^ were significantly upregulated, while those related to carbohydrate metabolism and translation were downregulated (Fig. 1c and Extended Data Fig. 2b,c and Supplementary Table 1,2). This comprehensive transcriptional analysis suggests that *Bb* persistence is impaired when preferred dietary fibers are not available as a carbon source, independent of any incumbent microbiota.

Next, we sought to identify common dietary fibers capable of restoring *Bb* persistence on the AIN+IP diet. We next tested raffinose, a plant-derived fiber widely found in legumes, vegetables, and whole grains. Since raffinose is a non-digestible trisaccharide degraded by α-galactosidase-producing microbes in the gut, raffinose may accumulate in the cecum of germ-free mice^38^. We further noted that a set of genes, *rafA* and *rafBCD*, potentially associated with raffinose metabolism and recently implicated in *B. breve* colonization of mono-colonized mice^39^, was also downregulated upon the diet switch (Fig. 1c). Furthermore, raffinose supported *in vitro* growth of *Bb* more effectively than other tested carbohydrates including HMOs (Extended Data Fig. 3a). When 6.5% (w/v) raffinose was added to the drinking water of *Bb* mono-colonized mice, the diet-dependent decrease in *Bb* colonization levels was fully rescued, reaching levels even higher than those on STD (Fig. 1d). Unlike inulin and pectin, 1% (w/v) raffinose sustained *Bb* colonization levels in mono-colonized mice on AIN diets (Fig. 1e), and significantly limited the expression of stress-response genes *groEL* and *ahpC* after the diet switch (Fig. 1f). Taken together, these data show that raffinose, a fiber directly utilized by *Bb*, can alleviate the diet switch-induced stress responses in intestinal *Bb,* and rescue *Bb* persistence in mono-colonized mice.

To expand this analysis beyond *Bb* to other common human-associated bifidobacteria, six additional *Bifidobacterium* species were tested to determine the effects of the diet switch and raffinose supplementation on their persistence in mono-colonized mice. Sensitivity to the diet switch was strictly conserved across species (Fig. 2a,b), suggesting that fermentable dietary fiber is critical for efficient intestinal colonization and persistence. Notably, *Bifidobacterium longum* subsp. *longum*, *Bifidobacterium longum* subsp. *infantis*, *Bifidobacterium animalis* subsp. *animalis*, and *Bifidobacterium animalis* subsp. *lactis* were efficiently rescued by raffinose, suggesting these four strains directly and efficiently utilize raffinose. By contrast, *Bifidobacterium bifidum* was not rescued by raffinose, and *Bifidobacterium adolescentis* showed a delayed and modest rescue effect (Fig. 2a,b). *B. bifidum* also showed severely impaired growth on raffinose as a sole carbon source *in vitro* (Extended Data Fig. 3b).

**Fig. 2.**
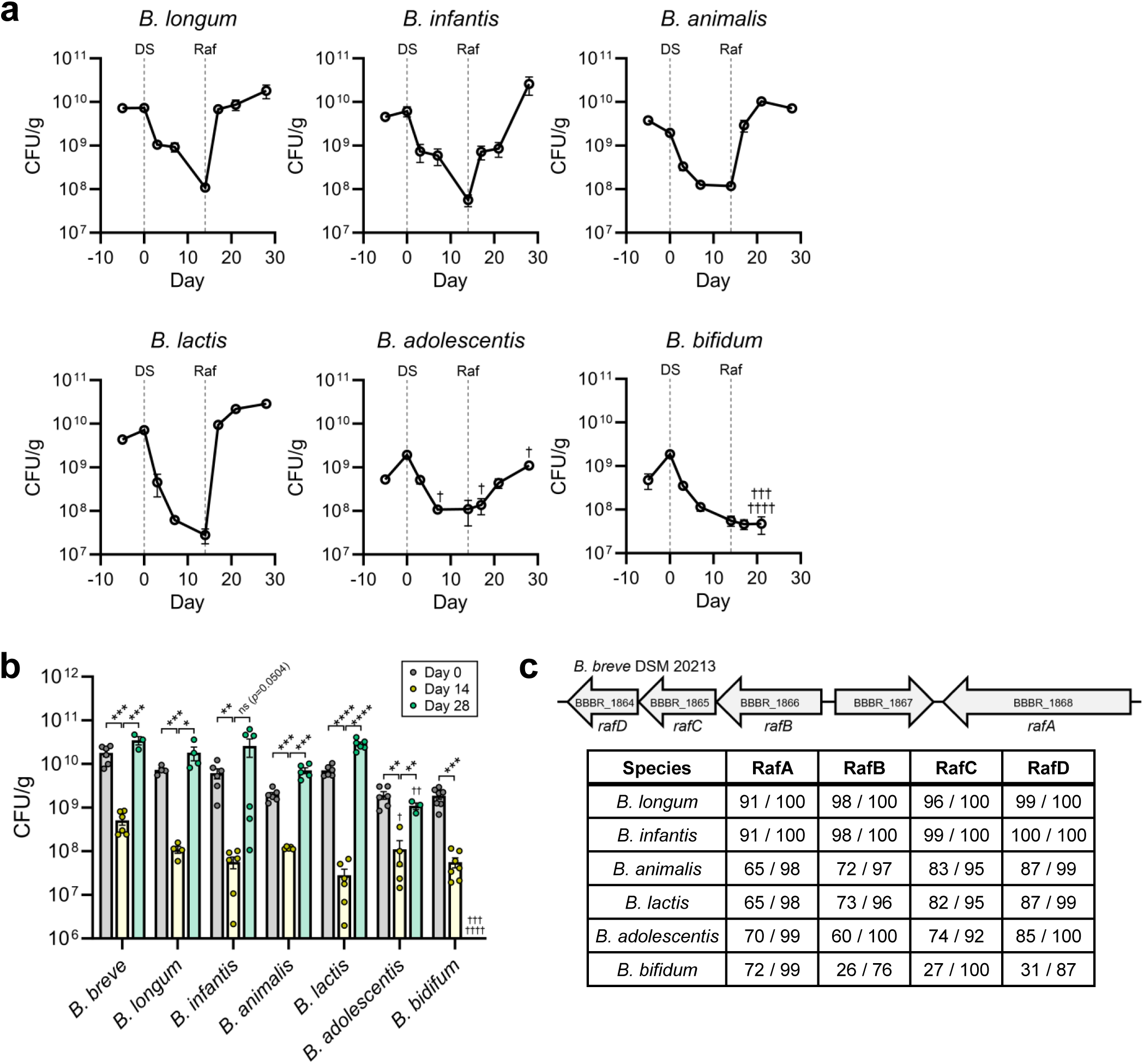
The *raf* operon confers raffinose exploitation by *Bifidobacterium* species. **a**,**b**, Germ-free mice were mono-colonized with the indicated *Bifidobacterium* species and maintained on STD for 10 days before being switched to AIN+IP. After 2 weeks on AIN+IP, mice were given drinking water supplemented with 6.5% (w/v) raffinose. Fecal CFUs of each species were quantified over time by plating. Data were pooled from two independent experiments (*n* = 4-5 for *B. longum*; *n* = 6 for *B. infantis*; *n* = 5-6 for *B. animalis*; *n* = 6 for *B. lactis*; *n* = 6 for *B. adolescentis*; *n* = 7 for *B. bifidum*). Mouse deaths are indicated with daggers. **b**, For statistical analysis, fecal CFUs of each species at day 0, 14 (the day raffinose was added to drinking water), and 28 (2 weeks after raffinose supplementation) after diet switch were analyzed. Data for *Bb* were obtained from Fig. 1c. Data are shown as mean ± SEM and were compared by Student’s *t*-test (*ns* not significant; \**p* < 0.05; \*\**p* < 0.005; \*\*\**p* < 0.0005; \*\*\*\**p* < 0.0001). **c**, A schematic of the *raf* operon in *Bb* is shown in the top panel. Amino acid sequence similarity of RafA-D proteins from the indicated *Bifidobacterium* species was assessed by BLASTP using the *Bb* proteins as references. Values in the table indicate % identity / % query coverage.

To explore the genetic basis of these species-specific raffinose responses, protein sequences of RafA-D were compared across species. RafA, an α-galactosidase, was broadly conserved across all species tested, including *B. bifidum* and *B. adolescentis* (Fig. 2c and Extended Data Fig. 4a). By contrast, *B. bifidum* showed low RafB-D sequence homology compared to raffinose-utilizing species (Fig. 2c), suggesting potentially impaired raffinose uptake. Although *B. adolescentis* retained high overall sequence similarity of RafB, a multiple sequence alignment identified amino acid residues that differed between both *B. bifidum* and *B. adolescentis* compared to the four efficient raffinose-utilizing species (Extended Data Fig. 4b). Structural alignment suggests that these variable residues may distort the predicted raffinose-binding pocket of RafB in *B. adolescentis* (Extended Data Fig. 4c). Given the putative role of RafB in competitive raffinose uptake^40^, these observations argue that species-specific variation in RafB-mediated raffinose transport contributes to differential raffinose responsiveness among *Bifidobacterium* species.

These data led us to hypothesize that efficient raffinose utilization confers a competitive advantage to *Bb* in the presence of other intestinal bacteria. To test this hypothesis, we assembled defined bacterial consortia that include up to 5 keystone species from the predominant human gut phyla (referred to as 5CON, 4CON, and 2CON; see methods). When *Bb* was co-inoculated with 5CON into germ-free mice, intestinal colonization levels of *Bb* were only slightly lower than those in mono-colonized mice under both STD and AIN+IP conditions (Fig. 3a). By contrast, other gnotobiotic bacterial consortia have been reported to severely limit or abolish bifidobacterial colonization^41,42^. Despite the nominal negative impact of 5CON on *Bb* colonization, a diet switch from STD to AIN+IP still significantly reduced *Bb* colonization in 5CON-colonized mice (Fig. 3a). To investigate how the diet switch affects *Bb+*5CON members, fecal microbiota composition was analyzed using 16S rRNA gene sequencing at day 0 and day 7 after the diet was switched from STD to AIN+IP. Strikingly, *Bb* was the only strain whose relative abundance was significantly decreased after the diet switch (Fig. 3b and Supplementary Table 3).

**Fig. 3.**
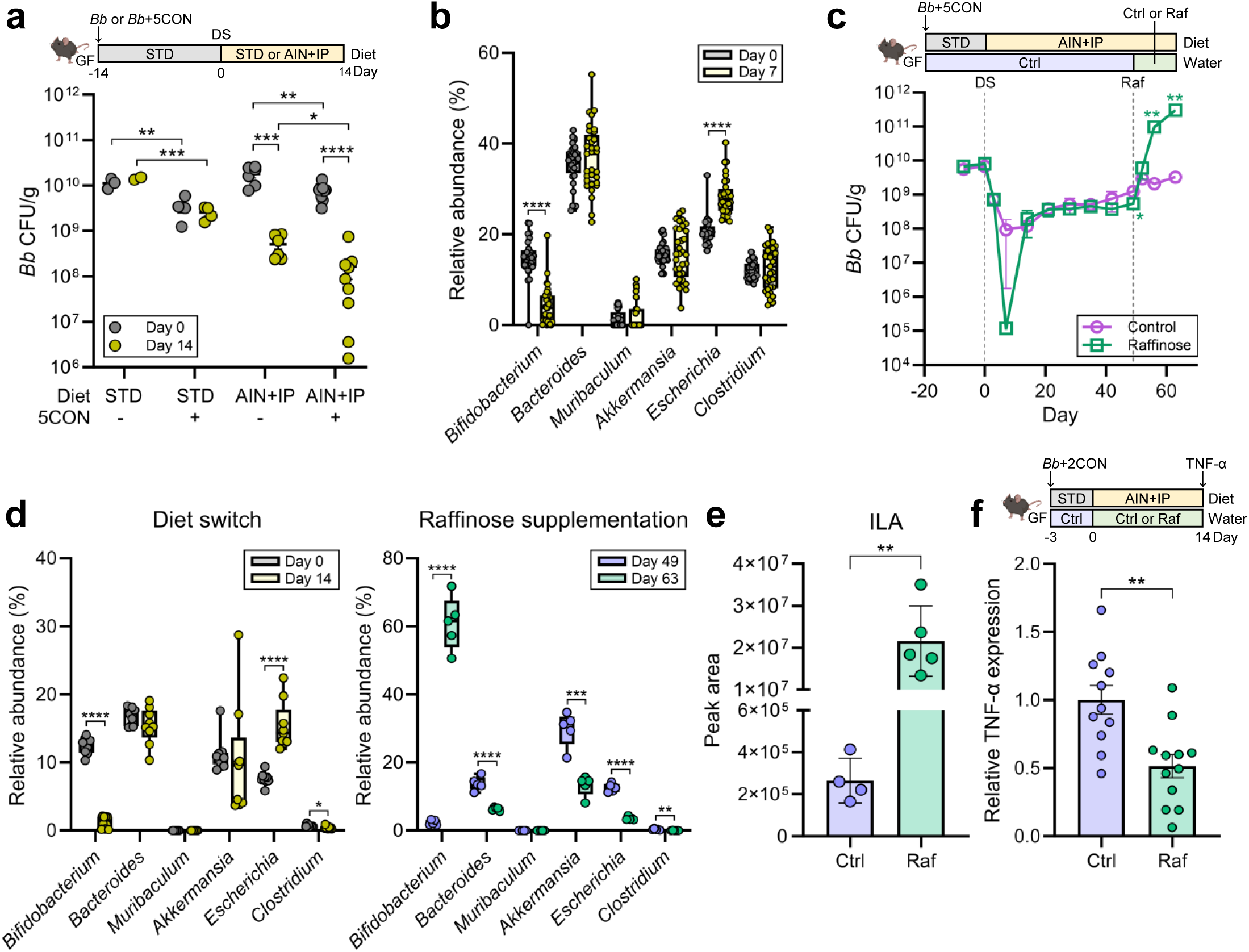
Diet and raffinose supplementation selectively modulate *Bb* colonization among common intestinal bacteria. **a**, Germ-free mice were colonized either with *Bb* alone or *Bb*+5CON and maintained on STD for 2 weeks. Then, mice were either continued on STD or switched to AIN+IP, as indicated by the diet label on the X-axis. Fecal *Bb* CFUs were quantified over time by plating. Data for the STD groups represent *n* = 2-3 for *Bb* and *n* = 4 for *Bb*+5CON. Data for the AIN+IP groups were pooled from two independent experiments (*n* = 6 for *Bb*; *n* = 9 for *Bb*+5CON). **b**, Germ-free mice were colonized with *Bb*+5CON and maintained on STD for 3 weeks before being switched to Fe-modified AIN+IP. Fecal microbiota at day 0 and day 7 after diet switch were analyzed by 16S rRNA gene sequencing. Both low-Fe and high-Fe AIN+IP were used and pooled for analysis because they showed comparable effects on fecal bacterial composition. Data were pooled from four independent experiments (*n* = 33). **c-e**, Germ-free mice were colonized with *Bb*+5CON, followed by a switch from STD to AIN+IP. After 7 weeks on AIN+IP, mice were given either control drinking water or drinking water supplemented with 6.5% (w/v) raffinose. Data represent *n* = 4 for control and *n* = 5 for raffinose. **c**, Fecal *Bb* CFUs were quantified over time by plating. **d**, Fecal microbiota at day 0 (day of diet switch), 14, 49 (day of raffinose supplementation), and 63 after diet switch were analyzed by species-specific qPCR. The relative abundance of each bacterium was normalized to total bacterial 16S rRNA levels. Data in **b**,**d** are shown as box-and-whisker plots with individual data points representing each mouse and were compared by Student’s *t*-test (\*\**p* < 0.005; \*\*\**p* < 0.0005; \*\*\*\**p* < 0.0001). **e**, Cecal indole-3-lactic acid (ILA) levels were measured by metabolomics at day 63. **f**, Germ-free mice were colonized with *Bb*+2CON and maintained on STD for 3 days. Then, mice were switched to AIN+IP with or without 4% (w/v) raffinose-supplemented water. Two weeks later, recombinant TNF-α (10 µg/mouse) was injected intraperitoneally to induce acute ileitis. Ileal TNF-α expression 2 hours after TNF-α injection was compared by qPCR and normalized to the control group. Data were pooled from three independent experiments (*n* = 11 for control; *n* = 12 for raffinose). Data in **a**,**c**,**e**,**f** are shown as mean ± SEM and were compared by Student’s *t*-test (\**p* < 0.05; \*\**p* < 0.005; \*\*\**p* < 0.0005; \*\*\*\**p* < 0.0001).

Consistent with the data in the mono-colonization model, supplemental raffinose in drinking water robustly and significantly rescued the diet-dependent decrease of *Bb* colonization in 5CON-colonized mice (Fig. 3c). qPCR analyses further showed that raffinose selectively increased the relative abundance of *Bb* (Fig. 3d), suggesting that both the diet switch and raffinose supplementation uniquely influence *Bb* in this model. Moreover, the increase in *Bb* colonization was raffinose dose-dependent in 4CON-colonized mice, and 4% raffinose in drinking water was sufficient to saturate intestinal colonization levels of *Bb* (Extended Data Fig. 5a). Finally, raffinose administration at the time of weaning still showed improved *Bb* persistence, demonstrating that this effect is not age-dependent (Extended Data Fig. 5b).

We next asked whether *Bb* provides raffinose-dependent host benefits in defined consortium models, as hypothesized based on prior studies. Cecal indole-3-lactic acid (ILA) levels were significantly increased in the raffinose-supplemented group compared to the control group (Fig. 3e). ILA is a beneficial metabolite produced at high levels by certain *Bifidobacterium* species (Extended Data Fig. 6a) and *Lactobacillus*^43^, and shows anti-inflammatory effects via AhR^11,44^. Given that *Bb* suppressed the expression of proinflammatory cytokines IL-6, IL-17, and IL-23 in the ileum of healthy 4CON-colonized mice (Extended Data Fig. 6b), we employed a TNF-driven inflammation model^45^ to examine whether raffinose mitigates acute ileal inflammation. Raffinose significantly reduced ileal Tnf-α expression 2 h after recombinant TNF injection (Fig. 3f), suggesting a dampened inflammatory feedback loop. Thus, raffinose supports *Bb* colonization specifically and selectively and provides anti-inflammatory effects associated with *Bb* in the defined consortium models. These observations raised the hypothesis that *Bb* may be uniquely able to utilize raffinose even in a more complex, physiologically relevant microbiota context.

To develop a physiologically-complex model of rapid microbiota diversification during human weaning, gnotobiotic mice initially colonized with *Bb*+2CON were conventionalized via fecal microbiota transplantation (FMT), using conventional mice purchased from Jackson Laboratories (Jax) as donors. When mice were switched to AIN at the time of Jax FMT, fecal *Bb* levels decreased to the limit of detection within 1-2 weeks (Fig. 4a), consistent with previous reports that bifidobacteria fail to stably colonize conventional mice. Remarkably, raffinose supplementation maintained robust *Bb* persistence after FMT (Fig. 4a). Given that responsiveness to dietary interventions can vary depending on microbiota composition, we repeated the experiment using fecal material from multiple independent donor colonies, including conventional SPF mice from another vendor (Charles River Laboratories; CRL) and an in-house SPF mouse colony (in-house), and a breastfed human infant fecal sample (Extended Data Fig. 7a). All tested FMT from different donors resulted in a dramatic loss of intestinal *Bifidobacterium* upon diet switch, which was fully rescued by raffinose supplementation (Fig. 4b and Extended Data Fig. 7b). This consistent response across four FMT donors implies that raffinose provides a selective advantage to raffinose-utilizing bifidobacteria within complex intestinal microbial communities. We further found that raffinose supported the expansion of endogenous *Bifidobacterium* in naïve SPF mice purchased from Jackson Laboratories, which was identified as *Bifidobacterium choerinum* by 16S rRNA gene sequencing (Extended Data Fig. 7c,d and Supplementary Table 4). By contrast, expansion of endogenous CRL *Bifidobacterium* driven by raffinose supplementation was more modest, reaching <1% relative abundance, in mice conventionalized using CRL FMT (Extended Data Fig. 7e), where the relative abundance of endogenous *Bifidobacterium* is lower than that in Jax FMT (Extended Data Fig. 7a). These results demonstrate that the effect of raffinose on bifidobacterial persistence is maintained in diverse complex microbiota, presumably because direct raffinose utilization confers a unique competitive advantage to certain bifidobacterial taxa.

**Fig. 4.**
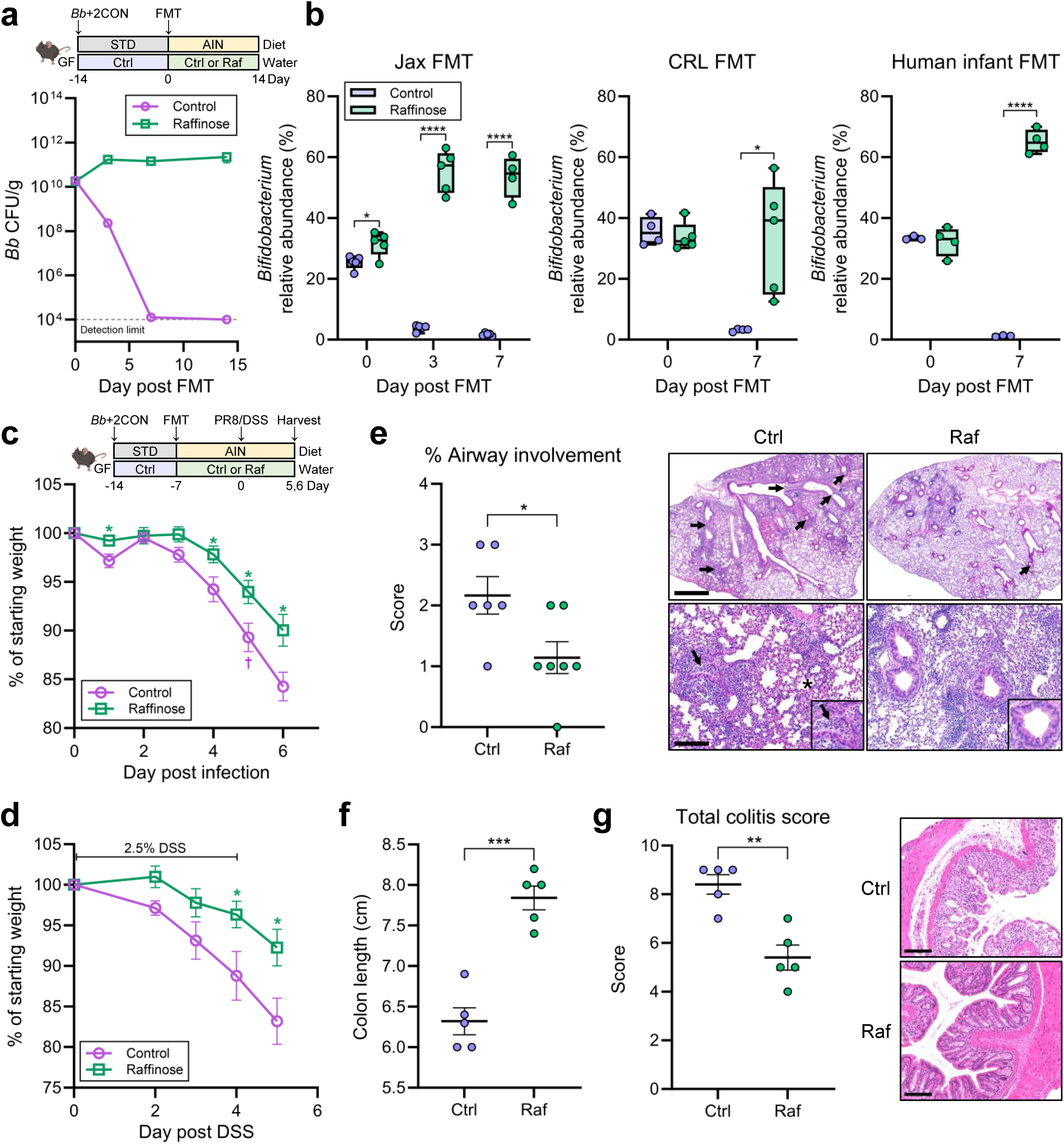
Raffinose supports *Bb* persistence and host health after microbiota diversification. **a**,**b**, Germ-free mice were colonized with *Bb*+2CON and maintained on STD for 2 weeks. Then, fecal microbiota from Jax or CRL SPF mice, or a human infant were introduced by oral gavage to induce microbiota diversification. Simultaneously, mice were switched to AIN with or without 6.5% (w/v) raffinose-supplemented water. Data represent *n* = 5 for Jax+Ctrl, *n* = 4-5 for Jax+Raf, *n* = 4 for CRL+Ctrl, *n* = 5 for CRL+Raf, *n* = 3 for human infant+Ctrl, and *n* = 4 for human infant+Raf. **a**, Fecal *Bb* CFUs in mice receiving Jax microbiota were quantified over time by plating. **b**, The relative abundance of *Bifidobacterium* after FMT and diet switch was analyzed by qPCR, using primers specific for the *Bifidobacterium* genus, and normalized to total bacterial 16S rRNA levels. Data are shown as box-and-whisker plots with individual data points representing each mouse. **c**-**g**, Germ-free mice were colonized with *Bb*+2CON and maintained on STD for one week. Then, mice received Jax FMT and were switched to AIN with or without 6.5% (w/v) raffinose-supplemented water. After an additional week, mice were either infected intranasally with PR8 (25 PFU; **c**,**e**) or administered 2.5% (w/v) DSS for 4 days (**d**,**f**,**g**). Disease progression was monitored by daily body weight changes relative to the starting weight (**c**,**d**). **e**, For the PR8 infection model, lungs were collected at day 6 post infection for hematoxylin and eosin (H&E) staining. Airway involvement was quantified as the percentage of affected airways in lung sections. Representative airways filled with inflammatory infiltrates and cellular debris are indicated by arrows and insets, and edema is marked by an asterisk. Scale bars, 1021 µm and 205 µm (top and bottom rows, respectively). For the DSS-induced colitis model, colon length (**f**) and distal colon histology (**g**) were analyzed at day 5 after DSS administration. **g**, Distal colon sections were stained with H&E and scored for inflammatory infiltration, epithelial damage, and tissue architecture disruption, with combined scores shown. Scale bars, 150 µm. Data in **c** were pooled from four independent experiments (*n* = 17 for control; *n* = 19 for raffinose). Data in **d** were pooled from two independent experiments (*n* = 9 for control; *n* = 10 for raffinose). Data in **e** (*n* = 6 for control; *n* = 7 for raffinose), **f** (*n* = 5 per group), **g** (*n* = 5 per group) are from one independent experiment. Data in **a**,**c**-**g** are shown as mean ± SEM and were compared by Student’s *t*-test (\**p* < 0.05; \*\*\**p* < 0.0005; \*\*\*\**p* < 0.0001).

We next asked whether this raffinose-driven *Bb* persistence after microbiota diversification could translate into functional benefits for the host. Given the established protective roles of *Bifidobacterium* species in influenza infection^13^ and DSS-induced colitis^46^, we tested whether raffinose improves disease outcomes in these models. To this end, mice colonized with *Bb*+2CON were introduced with Jax FMT and switched to AIN with or without raffinose. We confirmed that raffinose markedly promoted *Bb* levels, whereas levels in control mice dropped to near the detection limit by the time of A/Puerto Rico/8/1934 (PR8, H1N1) virus infection or DSS administration (Extended Data Fig. 8a,b). Notably, raffinose-fed mice exhibited significantly attenuated weight loss compared to controls in both disease models (Fig. 4c,d). Furthermore, in the PR8 infection model, raffinose-fed mice showed reduced airway involvement in lung tissue (Fig. 4e and Extended Data Fig 8c) and improved survival (Extended Data Fig. 8d). Similarly, in the DSS-induced colitis model, raffinose-fed mice exhibited reduced colon shortening and improved distal colon histopathology (Fig. 4f,g and Extended Data Fig. 8e). Collectively, these results suggest that raffinose contributes to improved host outcomes after microbiota diversification, in parallel with enhanced bifidobacterial persistence.

Finally, to gain mechanistic insight into raffinose-driven competitive persistence of bifidobacteria within a complex microbiota, we evaluated wild-type *B. breve* strain UCC2003 (WT) and isogenic Δ*rafA* and Δ*rafB* mutants^39^. In mono-colonized mice, all strains showed comparable decreases in colonization after the diet switch (Extended Data Fig. 9a). However, raffinose restored colonization of WT and Δ*rafB*, but not Δ*rafA* (Extended Data Fig. 9a). These data suggest RafA is strictly required for raffinose utilization *in vivo.* By contrast, RafB is not strictly required for raffinose utilization in the absence of other bacteria. In the Jax FMT diversification model, raffinose still promoted total *Bifidobacterium* levels, since *Bifidobacterium* was rapidly eliminated within three days of the switch to AIN diet in the absence of raffinose (Extended Data Fig. 9b). WT persisted for 3 days in the presence of raffinose and was subsequently replaced by the Jax-endogenous *Bifidobacterium* (Extended Data Fig. 9b,c). Meanwhile, although Δ*rafB* was promoted by raffinose in mono-colonized mice (Extended Data Fig. 9a), it was lost more rapidly than WT in the presence of raffinose after FMT (Extended Data Fig. 9c). These data illustrate a critical role for RafB-mediated raffinose transport in the competitive persistence of bifidobacteria within a diversified gut microbiota.

In this study, we demonstrated that dietary raffinose promotes robust bifidobacterial persistence in the mouse intestine across multiple dynamic diet and microbiome conditions, and confers protective effects in influenza infection and DSS-induced colitis in adult mice. Dietary raffinose provides a competitive advantage for bifidobacterial persistence, similar to well-studied HMOs^17,19,20,31^. These findings are consistent with recent reports of geographic variation in both bifidobacterial colonization levels and local dietary adaptations among bifidobacteria^16,47^, highlighting the critical role of specific dietary fibers in bifidobacterial persistence. Given its impact on host outcomes in both infectious and inflammatory disease models, dietary guidelines promoting consistent raffinose intake starting at weaning are a potential non-invasive strategy to improve host resilience and limit disease burden at the population level. Analogous evidence-based dietary guidelines for pediatric peanut intake have correlated with decreased food allergy rates at the population level^48,49^. The efficacy of dietary raffinose interventions may depend on the presence, or supplementation, of efficient raffinose-utilizing taxa, highlighting the potential for microbiome-directed precision nutrition approaches.

## Methods

### Mice

Germ-free and conventional SPF C57BL/6N mice were initially purchased from Charles River Laboratories, and germ-free *Rag1*^-/-^ mice were kindly shared by Dr. Gregory Sonnenberg. All gnotobiotic mice for this study were bred and maintained in the gnotobiotic core facility at Rutgers New Jersey Medical School under a 12h light/dark cycle. SPF C57BL/6J mice were purchased from Jackson Laboratories and then bred and maintained in the SPF facility at Rutgers New Jersey Medical School. Both male and female mice were used for experiments at 7-12 weeks of age unless otherwise indicated. To examine whether age-dependent effects of diet switch and raffinose supplementation on *Bb* colonization, pups from *Bb*+2CON-colonized dams were weaned and used at postnatal day 20. All animal experiments were performed according to the protocol approved by the Institutional Animal Care and Use Committee (IACUC) at Rutgers University.

### Diets and drinking water

Gnotobiotic mice were fed an autoclaved standard lab diet (LabDiet 6F 5KAI) and autoclaved water *ad libitum* unless otherwise noted. For diet-switch experiments, mice were moved into a fresh cage with free access to an irradiated AIN-93G-based diet (Teklad)^35^. To assess the effect of dietary iron on *Bb* colonization, Fe-modified AIN-93G diets were used, including a low-iron diet (3-6 ppm Fe) and a high-Fe diet (2,000 ppm Fe). Unless otherwise specified, the AIN-93G-based diet for gnotobiotic mice contained 35 ppm Fe, which is sufficient normal rodent growth and development, and was supplemented with 1% inulin and 1% pectin (AIN-IP). To avoid cross-feeding interactions within the microbiota, inulin and pectin were not supplemented to the AIN-93G diet (AIN) for conventionalized mice. To supplement raffinose in drinking water, D-(+)-raffinose pentahydrate (Millipore) was dissolved in ultra-purified water at the concentrations indicated in the corresponding figure legends and filtered through a 0.2 µm membrane for sterilization. Filtered ultra-purified water was used as a control^50^. Mice had free access to either control or raffinose-supplemented water during the experiments.

### Bacterial strains and culture conditions

All bacterial strains used in this study are listed in Supplementary Table 5. *Bifidobacterium* species and *Bacteroides thetaiotaomicron*, *Muribaculum intestinale*, *Akkermansia muciniphila*, and *Clostridium sporogenes* were obtained from ATCC or DSMZ. *Escherichia coli* Nissle 1917, packaged as ‘Mutaflor’, was purchased from feelgoodnatural.com. *B. breve* UCC2003 WT, Δ*rafA*, and Δ*rafB* strains were kindly provided by the van Sinderen laboratory^39^. For colonization of germ-free and gnotobiotic mice, bacterial overnight cultures were prepared by inoculating each strain into the appropriate culture medium (see Supplementary Table 5 for details) and incubating at 37℃ for 24-48 h under anaerobic conditions. Cultures were then combined to generate 200 µl inocula containing ∼10^8^ CFU of each bacterium for oral gavage. 5CON consisted of *B. thetaiotaomicron*, *M. intestinale*, *A. muciniphila*, *E. coli*, and *C. sporogenes*, whereas 4CON excluded *M. intestinale* due to poor colonization. 2CON consisted of *B*. *thetaiotaomicron* and *C. sporogenes* to avoid any potential confounding probiotic effects of *E. coli* Nissle 1917 and *A. muciniphila*, and also to more closely mimic the microbiome at weaning, where these two bacteria are rare.

### Fecal *Bifidobacterium* CFU enumeration

To quantify intestinal colonization levels of *Bifidobacterium* species, fresh fecal samples were collected and processed for CFU enumeration. Samples were weighed, homogenized in sterile phosphate-buffered saline (PBS), and serially diluted. For samples obtained from mono-colonized mice, dilutions were plated on MRS agar. For samples from consortium-colonized or conventionalized mice, dilutions were plated on MRS agar supplemented with 50 mg/L mupirocin (Sigma), which selectively permits the growth of *Bifidobacterium* species. Plates were incubated anaerobically at 37°C for 48 h prior to colony counting. In experiments involving *B. bifidum*- or Δ*rafA*-mono-colonized mice, severe diarrhea and weight loss were observed following raffinose supplementation, and the experiments were terminated at day 7 post supplementation.

### DNA and RNA isolation

To analyze bacterial composition and gene expression, total DNA and RNA were extracted from fecal samples using the QIAamp PowerFecal Pro DNA Kit or the AllPrep PowerFecal Pro DNA/RNA Kits (Qiagen), according to the manufacturer’s instructions. For RNA isolation, an additional DNase treatment step was included using RNase-Free DNase (Qiagen) to remove residual genomic DNA.

For host gene expression analysis, mouse ileal tissues were collected into Soft Tissue Lysing Tubes for Precellys24 (VWR) containing TRIzol™ Reagent (Invitrogen) and homogenized using a Precellys24 tissue homogenizer (Bertin Technologies). Total RNA was then isolated using the Direct-zol RNA Microprep Kit (Zymo Research) according to the manufacturer’s instructions.

### Bacterial RNA-seq and analysis

Total RNA extracted from fecal samples of *Bb* mono-colonized mice was used for bacterial RNA-seq. Bacterial rRNA was depleted from total RNA extracts using the NEBNext rRNA Depletion Kit (Bacteria) according to the manufacturer’s protocol. Briefly, 100 ng of total RNA was hybridized with single-stranded DNA probes that bind to ribosomal RNA. The resulting RNA:DNA hybrids were selectively degraded via RNase H digestion, and the remaining DNA probes were removed using DNase I treatment. The rRNA-depleted RNA was then purified using Agencourt RNAClean XP beads (Beckman Coulter).

Subsequently, the depleted RNA was fragmented and reverse transcribed with random primers using the NEBNext Ultra II RNA Library Prep Kit. Double-stranded cDNA was generated using the NEBNext Second Strand Synthesis Enzyme Mix for 1 hour at 16°C. The cDNA was then end-repaired and dA-tailed, followed by adaptor ligation. Adaptor-ligated fragments were purified using AMPure XP beads and enriched by PCR amplification using NEBNext Ultra II Q5 Master Mix and indexed primers.

The final libraries were purified using AMPure XP beads, and library size distribution was assessed using D1000 ScreenTape on an Agilent TapeStation system. Libraries were quantified using a Qubit fluorometer (Thermo Fisher Scientific), pooled in equimolar ratios, and sequenced on an Illumina NextSeq 6000 instrument (Illumina) in a single-end 1 × 100 bp configuration.

The raw sequencing reads were aligned to *Bb* (*B. breve* DSM 20213) reference genome (NCBI accession number: AP012324) using STAR (v2.7.11b)^51^, and gene-level read counts were generated using HTSeq-count (v2.0.7)^52^. Differential expression analysis and principal component analysis (PCA) were performed in R (v4.3.1) using DESeq2 (v1.42.1)^53^, using only genes having 10 or more counts in at least four samples. Log2 fold changes were calculated using DESeq2 by comparing day 2, day 4, and day 35 samples to the day 0 reference condition. PCA results were visualized using ggplot2 and ggforce in R. Differential expression results were visualized as heatmaps and volcano plots in GraphPad Prism and further used to run gene set enrichment analysis (GSEA). GSEA was run in R (v4.1.0) using clusterProfiler (v4.2.2). The bitr_kegg command was used to convert gene IDs via the Kyoto Encyclopedia of Genes and Genomes (KEGG) API, and the gseKEGG function was used to run GSEA using KEGG.

### Quantitative PCR (qPCR)

Relative abundance of each bacterium was determined by qPCR using total DNA extracted from fecal samples and bacteria-specific primers (Supplementary Table 6). For bacterial gene expression analysis, complementary DNA (cDNA) was synthesized from total RNA extracted from fecal samples using the iScript™ cDNA Synthesis Kit (Bio-Rad), according to the manufacturer’s instructions. For host gene expression analysis, cDNA was synthesized from ileal tissue RNA using the SuperScript™ First-Strand Synthesis System (Invitrogen), following the manufacturer’s protocol. qPCR was performed using PowerUp™ SYBR™ Green Master Mix (Applied Biosystems) on a CFX Opus 384 Real-Time PCR System (Bio-Rad). Bacterial and host gene expression levels were normalized to *rnpA*^54^ and *Hprt*^55^, respectively, as internal controls. Relative gene expression levels were calculated using standard normalization methods. All gene-specific primers used in this study are listed in Supplementary Table 6. Specificity of qPCR primers was confirmed by control experiments with genomic DNA from pure cultures of 5CON bacteria and *Bifidobacterium* strains.

### Protein sequence and structural alignments

Amino acid sequences of RafA-D proteins encoded in the *raf* operon from *Bb* (*B. breve* DSM 20213) were used as queries in NCBI BLASTP searches^56^ to identify homologs from the indicated *Bifidobacterium* species. For each query, the top-scoring hit was selected, and the percentage identity and query coverage were extracted from the alignments. Selected RafA and RafB homolog sequences were then aligned using default settings in the Clustal Omega portal (https://www.ebi.ac.uk/jdispatcher/msa/clustalo)^57^. For structural alignments, PDB 4ZS9, containing RafB from *B. lactis* Bl-04 bound to raffinose^40^, was globally aligned with an AlphaFold-modeled structure of the closest RafB homolog in *B. adolescentis* ATCC 15703 (https://alphafold.com/entry/AF-A1A3S2-F1)^58^.

### *In vitro* growth kinetics assay

*In vitro* growth kinetics of *Bb* and *B. bifidum* were evaluated in carbohydrate-defined media. Growth media were prepared using Lactobacillus MRS broth without dextrose (Alpha Biosciences) supplemented with 0.05% (w/v) L-cysteine and 2% (w/v) of the indicated carbohydrate, including glucose (Sigma), raffinose, lactulose (Sigma), 2′-fucosyllactose (2′FL; Glycom), lacto-*N*-tetraose (LNT; Glycom), or lacto-*N*-neotetraose (LNnT; Glycom), as the sole carbon source. Overnight cultures were grown anaerobically at 37°C for 24-48 h and inoculated into 96-well plates at an initial OD_600_ of 0.1 in a final volume of 200 µl per well. Plates were then sealed with MicroAmp™ Optical Adhesive Film (Applied Biosystems) and incubated at 37°C with shaking in a Varioskan LUX plate reader (Thermo Fisher Scientific). OD_600_ was measured every 30 min for 20 h (*Bb*) or 24 h (*B. bifidum*). Blank-subtracted values were used for analysis. Data represent the mean of technical triplicates from three independent experiments.

### 16S rRNA gene sequencing

To characterize microbial composition, the V3–V4 variable regions of the 16S rRNA gene were amplified from fecal DNA by PCR. Briefly, 20 ng of fecal DNA was used to generate amplicons using an Illumina-compatible library preparation protocol. The V3–V4 regions were amplified using locus-specific primers with Illumina overhang adapter sequences (Supplementary Table 6). PCR products were purified using Agencourt AMPure XP beads (Beckman Coulter). A second PCR was performed using the purified amplicons as templates to attach dual indices and Illumina sequencing adapters with the Nextera XT Index Kit (Illumina). The quality and quantity of the enriched indexed libraries were assessed using an Agilent TapeStation and Qubit fluorometer, respectively. Libraries were then pooled at equimolar concentrations and sequenced on the Illumina MiSeq platform (Illumina) using the MiSeq v3 reagent kit (2 × 300 bp paired-end reads).

Raw paired-end reads were assessed and trimmed on quality with FastQC (v0.11.9) and cutadapt (v3.4)^59^, respectively. Amplicon sequence variants (ASVs) were inferred using DADA2 (v1.22.0)^60^ and assigned taxonomy with a Naive Bayes classifier to the Silva (v138) database^61,62^. ASVs were agglomerated to the genus level for subsequent downstream analyses.

### Metabolite extraction

Cecal metabolites were extracted with methanol. 30 mg of dried cecal content was vortexed and sonicated in 500 μl of 100% methanol for 1 min. Samples were vortexed and sonicated for 1 min. Cecal extracts were pelleted at 5,000 *g* for 10 min at 4℃, and the supernatants were transferred to 2 ml HPLC vials. Samples were then dried in a SpeedVac (Thermo Fisher Scientific) vacuum concentrator. Dried materials were resuspended in 1 ml of methanol and vortexed for 1 min. The samples were pelleted at 5,000 *g* for 5 min at 22℃, and the supernatants were transferred to HPLC vials and dried in a SpeedVac vacuum concentrator. The samples were then resuspended in 100 μl of methanol and centrifuged at 5,000 *g* for 10 min at 22℃. Clarified extracts were transferred to fresh HPLC vials and stored at -20℃ until analysis.

### MS analysis

LC−MS was performed on the Thermo Fisher Scientific Vanquish UHPLC system coupled with a Thermo Q Exactive HF hybrid quadrupole-orbitrap high-resolution mass spectrometer equipped with a HESI ion source. Metabolites were separated using a water−acetonitrile gradient on an Agilent Zorbax Eclipse XDB-C18 column (150 mm × 2.1 mm, particle size 1.8 μm) maintained at 40℃ (solvent A: 0.1% formic acid in water; solvent B: 0.1% formic acid in acetonitrile). The A/B gradient started at 1% B for 3 min after injection and increased linearly to 100% B at 20 min, then 100% B for 5 min, and down to 1% B for 3 min at a flow rate of 0.5 ml min^−1^. MS parameters were as follows: spray voltage, 3.5 kV; capillary temperature, 380℃; probe heater temperature, 400℃; 60 sheath flow rate, 20 auxiliary flow rate, and one spare gas; S-lens RF level, 50; resolution, 240,000; AGC target, 3 × 10^6^. The instrument was calibrated weekly with positive and negative ion calibration solutions (Thermo Fisher Scientific). Each sample was analyzed in negative and positive ionization modes using an *m*/*z* range of 100 to 800. Data were collected using Thermo Fisher Scientific Xcalibur software (v4.1.31.9) and quantified by integration in Excalibur Quan Browser (v4.1.31.9).

### Feature detection and characterization

LC−MS RAW files from biospecimens from control and raffinose-supplemented mice, or control and *Bb-*colonized mice, were converted to mzXML format (centroid mode) using MSconvert (v3.0.20315-7da487568; ProteoWizard), followed by analysis using the XCMS analysis feature in Metaboseek (v0.9.7; https://metaboseek.com)^63^ based on the centWave XCMS algorithm to extract features^64^. Peak detection values were set as follows: 4 ppm, 3 to 20 peakwidth, 3 snthresh, 3 and 100 prefilter, FALSE fitgauss, 1 integrate, TRUE firstBaselineCheck, 0 noise, wMean mzCenterFun and −0.005 mzdiff. The identity of ILA, was confirmed by injection with an authentic standard, which confirmed the retention time of ILA as 8.10435 minutes, with an *m*/*z* value of 204.06671 for [M-H]^-^ detected in negative ionization mode.

### TNF-α-induced acute ileitis model

Germ-free mice were colonized with *Bb*+2CON by oral gavage. Three days after colonization, mice were switched to AIN+IP with or without 4% (w/v) raffinose in drinking water and maintained for 2 weeks. Then, acute ileitis was induced by intraperitoneal injection of recombinant mouse TNF-α (10 µg in 200 µl PBS; BioLegend). Mice were euthanized 2 h post-injection, and ileal tissues were collected for gene expression analysis.

### Conventionalization of gnotobiotic mice by FMT

FMT material was prepared from conventional SPF mice obtained from Jackson Laboratories and Charles River Laboratories, as well as from in-house bred SPF mice maintained in the SPF facility at Rutgers New Jersey Medical School. Cecal and colonic contents were collected, resuspended in sterile PBS, and the supernatant was aliquoted and stored at -80℃ until use. A human infant fecal sample was kindly provided by the Domínguez-Bello laboratory^65^ and processed similarly by resuspension in PBS, followed by collection of the supernatant and storage at -80℃. To conventionalize gnotobiotic mice, 200 µl of thawed FMT aliquots were administered by oral gavage. Mice were subsequently transferred from the gnotobiotic core facility to the SPF facility following FMT.

### Influenza infection model

Germ-free mice were colonized with *Bb*+2CON and maintained on STD. After a week of colonization, mice were conventionalized by oral gavage with Jax FMT. At the time of FMT, mice were switched to AIN and provided drinking water supplemented with or without 6.5% (w/v) raffinose. A week after conventionalization and diet switch, at day 0, mice were anesthetized with ketamine–xylazine, and intranasally infected with 25 plaque-forming units (PFU) of PR8 (A/Puerto Rico/8/1934; H1N1) virus in 50 µl of serum-free DMEM. Fecal samples were collected at the time of FMT and at the day of infection to count *Bb* CFUs. Disease progression was monitored by measuring body weight daily at a consistent time point and expressed relative to the initial body weight at the day of infection. Humane endpoints were defined as a loss of ≥25% of initial body weight on day 0. Survival was monitored for up to 17 days post-infection. For histological analysis, mice were anesthetized intraperitoneally by ketamine-xylazine, and lungs were transcardially perfused with 10 ml PBS. The lung tissues were then collected and fixed in 4% paraformaldehyde for 24 hours. Lung tissues stained with hematoxylin and eosin (H&E) were examined by a boarded pathologist using the post-examination method of evaluation to enhance reproducibility and prevent bias^66,67^. Lung tissues were evaluated for airway obstruction by inflammation and cellular debris using ordinal scores: 0, none; 1, <25%; 2, <50%; 3, <75%; and 4, >75% of affected airways in tissue section. Pulmonary edema in airspaces^68^, a sign of acute lung injury, was also evaluated: 0, none; 1, <25%; 2, <50%; 3, <75%; and 4, >75% of 200x fields affected.

### DSS-induced colitis model

Conventionalized mice were prepared as described above for the influenza infection model. Briefly, age-matched germ-free male mice colonized with *Bb*+2CON were conventionalized with Jax FMT and switched to AIN with or without 6.5% (w/v) raffinose in drinking water. At day 0, colitis-grade DSS (average MW 36,000-50,000 Da; MP Biomedicals) was added to the drinking water at a concentration of 2.5% (w/v). Fecal samples were collected at the time of FMT and at the day of DSS administration to count *Bb* CFUs. DSS was administered *ad libitum* for 4 days to induce significant inflammation, as indicated by ∼10% weight loss, after which DSS was removed from the drinking water. The body weight of mice was measured daily at a consistent time point and expressed relative to the initial body weight at the day of DSS administration. At day 5 post-DSS administration, colon tissues were collected for length measurement and histological analysis. Distal colon tissues were fixed in 4% paraformaldehyde for 48 hours and stained with H&E. Slides were scored on three different factors: 1.) inflammatory infiltration (0-3), 2.) epithelial damage (0-3), 3.) tissue architecture disruption (0-3)^69^. Scores for the three factors were combined into a total colitis score.

## Supporting information

Supplementary Table 1 DEG analysis of fecal Bb after diet switch

Supplementary Table 2 GSEA of fecal Bb after diet switch

Supplementary Table 3 Genus-level microbiota composition in Bb+5CON mice after diet switch

Supplementary Table 4 Genus-level microbiota composition in conventional Jax mice after diet switch and raffinose supplementation

Supplementary Table 5 Bacterial strains used in this study

Supplementary Table 6 Primers used in this study

## Data availability

The bacterial RNA-seq and 16S rRNA gene sequencing datasets are available in GEO under accession numbers GSE330259 for bacterial RNA-seq, and GSE330841 and GSE330838 for 16S rRNA gene sequencing.

## Acknowledgements

We thank members of the Bessman laboratory for critical discussions and feedback on this manuscript, the Rutgers NJMS Cellular Imaging and Histology core for technical assistance, Dr. Gregory Sonnenberg for germ-free *Rag1*^-/-^ mice, and DSM-Firmenich for HMO reagents. This work was supported by grants from the National Institutes of Health to N.J.B. (DK128308 and AI154440), L.Y.R.W. (R00AI170996), F.C.S. (R35 GM131877), a grant from Science Foundation Ireland (SFI; now ResearchIreland) to D.V.S. (12/RC/2273), and D.K.M. (P30 DK054759, Comparative Pathology Core in the Precision Medicine Center for Cystic Fibrosis)

## Contributions

Conceptualization: G.C. and N.J.B. Methodology: G.C., S.C., T.H.W., H.S., A.L., D.K.M., M.G.D, F.C.S., D.v.S., L.Y.R.W., and N.J.B. Investigation: G.C. performed most experiments, with assistance from S.C., U.S., A.S., J.K., and W.M. Influenza experiments were performed collaboratively with G.C., S.C., and L.Y.R.W. Metabolomics was performed and analyzed by T.H.W., M.A.F., and F.C.S. Infant fecal samples were collected and processed by H.S. and M.G.D. D.v.S. developed genetic tools for bifidobacteria. G.C., A.M., and A.L. analyzed sequencing data.

## Extended Data Figures

**Extended Data Fig. 1.**
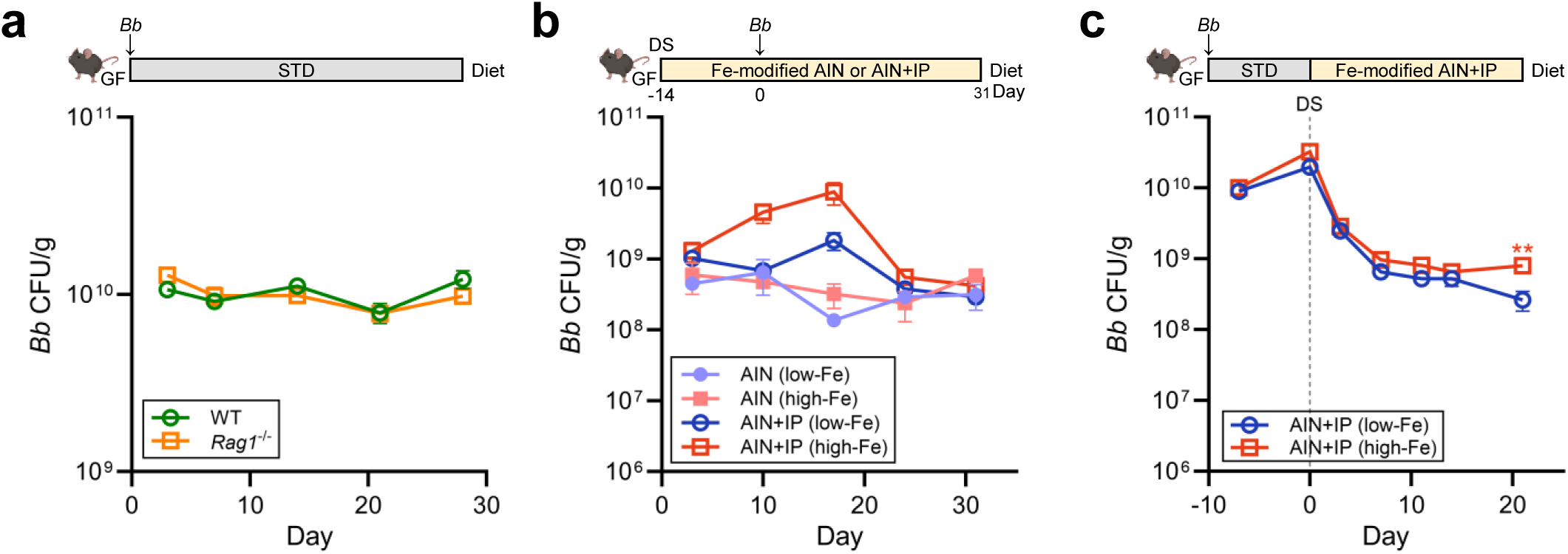
The host adaptive immune system and dietary iron do not impact *Bb* colonization in mono-colonized mice. **a**, Germ-free wild-type (WT) or *Rag1*^-/-^ mice were mono-colonized with *Bb* and maintained on STD throughout the experiment. Fecal *Bb* CFUs were quantified over time by plating. Data represent *n* = 5 for WT and *n* = 4-5 for *Rag1*^-/-^. **b**, Germ-free mice were switched from STD to Fe-modified AIN or Fe-modified AIN+IP. Two weeks later, mice were mono-colonized with *Bb,* and fecal *Bb* CFUs were quantified over time by plating. Data were pooled from one or two independent experiments (*n* = 3 for low-Fe AIN; *n* = 4 for high-Fe AIN; *n* = 10 for low-Fe AIN+IP; *n* = 10 for high-Fe AIN+IP). **c**, Germ-free mice were mono-colonized with *Bb* and maintained on STD for 10 days. Then, mice were switched to Fe-modified AIN+IP, and fecal *Bb* CFUs were quantified over time by plating. Data were pooled from two independent experiments (*n* = 9 for low-Fe AIN+IP; *n* = 10 for high-Fe AIN+IP). All data are shown as mean ± SEM and were compared by Student’s *t*-test (\*\**p* < 0.005).

**Extended Data Fig. 2.**
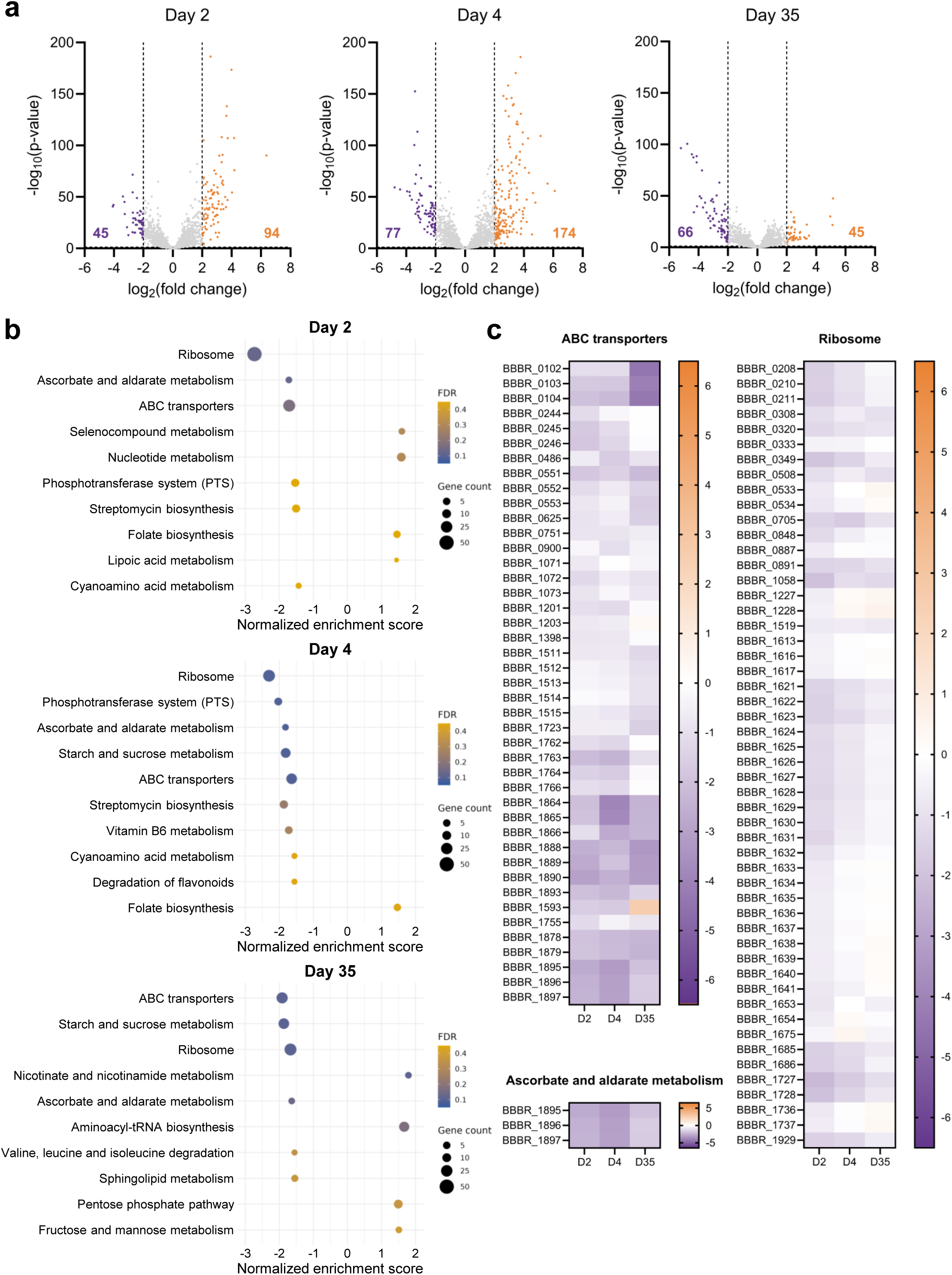
Diet switch induces a clear transcriptome shift of *Bb* in mono-colonized mice. Germ-free mice were mono-colonized with *Bb*, followed by a switch from STD to Fe-modified AIN+IP, as depicted in Fig. 1A. Both low-Fe and high-Fe AIN+IP were used and pooled for analysis because they showed comparable effects on *Bb* transcriptomes. Transcriptomic changes in *Bb* at day 0, 2, 4, and 35 after diet switch were analyzed by RNA-seq. Data were pooled from two independent experiments (*n* = 19). **a**, Volcano plots depicting differentially expressed genes (DEGs) following diet switch. DEGs were defined as genes with |log2(fold change)| ≥ 2 and adjusted *p* < 0.05. Orange and purple numbers denote the number of up-regulated and down-regulated genes, respectively. **b**, Pathway enrichment analysis of DEGs identified in (**a**). Gene set enrichment analysis (GSEA) was performed to assess pathway-level changes at day 2, 4, and 35 relative to day 0 after diet switch. The top 10 pathways ranked by false discovery rate (FDR)-adjusted *p*-value (Benjamini-Hochberg correction) are shown with normalized enrichment scores (NES). Pathways with an FDR-adjusted *p* < 0.1 were considered significantly enriched. **c**, Heatmap visualizing the relative expression of genes belonging to significantly altered pathways identified in (**b**). Gene expression levels at day 2, 4, and 35 were normalized to day 0.

**Extended Data Fig. 3.**
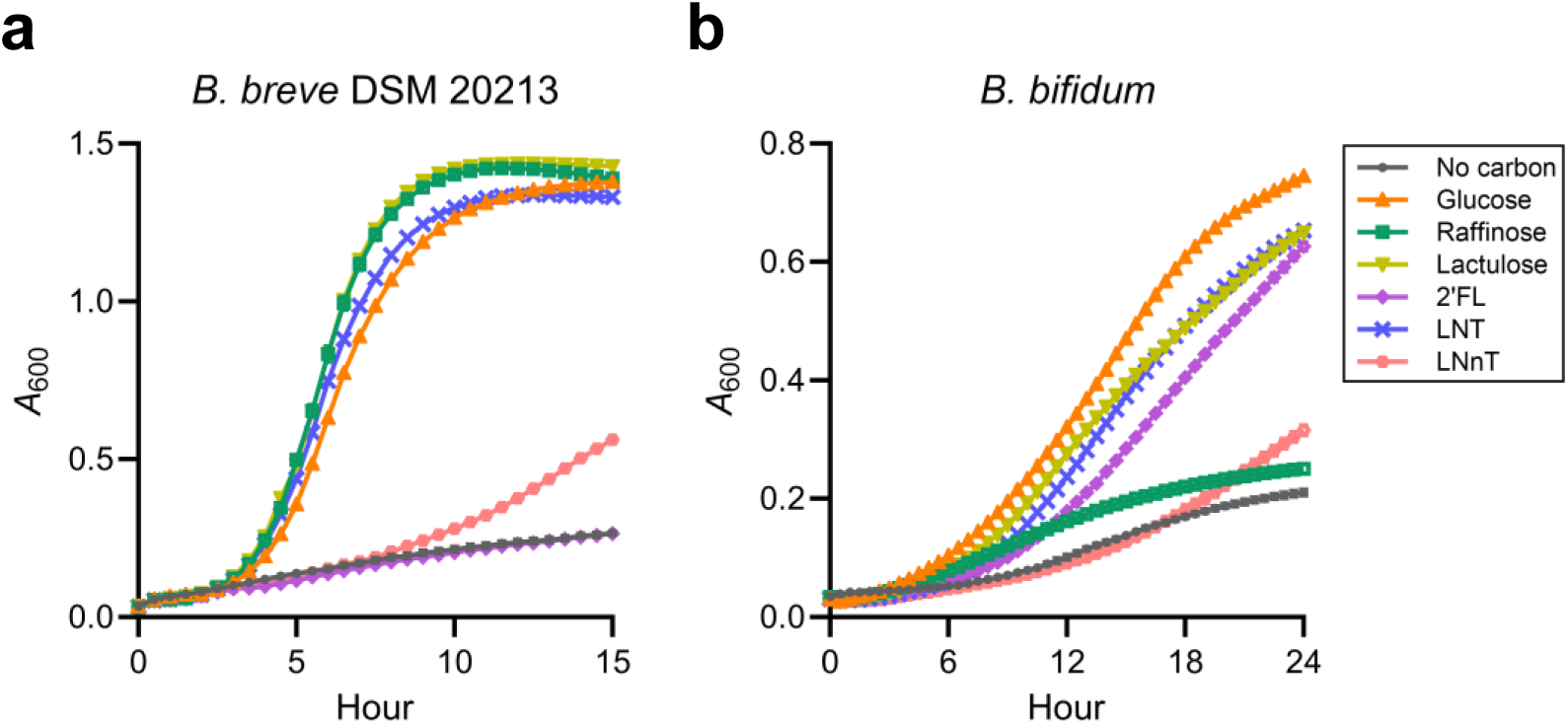
Raffinose-driven growth of *Bifidobacterium* is dependent on the presence of the functional *raf* operon. **a**,**b**, Growth curves of *Bb* (*B. breve* DSM 20213; **a**) and *B. bifidum* DSM 20456 (**b**) in dextrose-devoid MRS supplemented with 2% (w/v) of the indicated carbohydrates as the sole carbon source. All data were pooled from three independent experiments and are shown as mean ± SEM.

**Extended Data Fig. 4.**
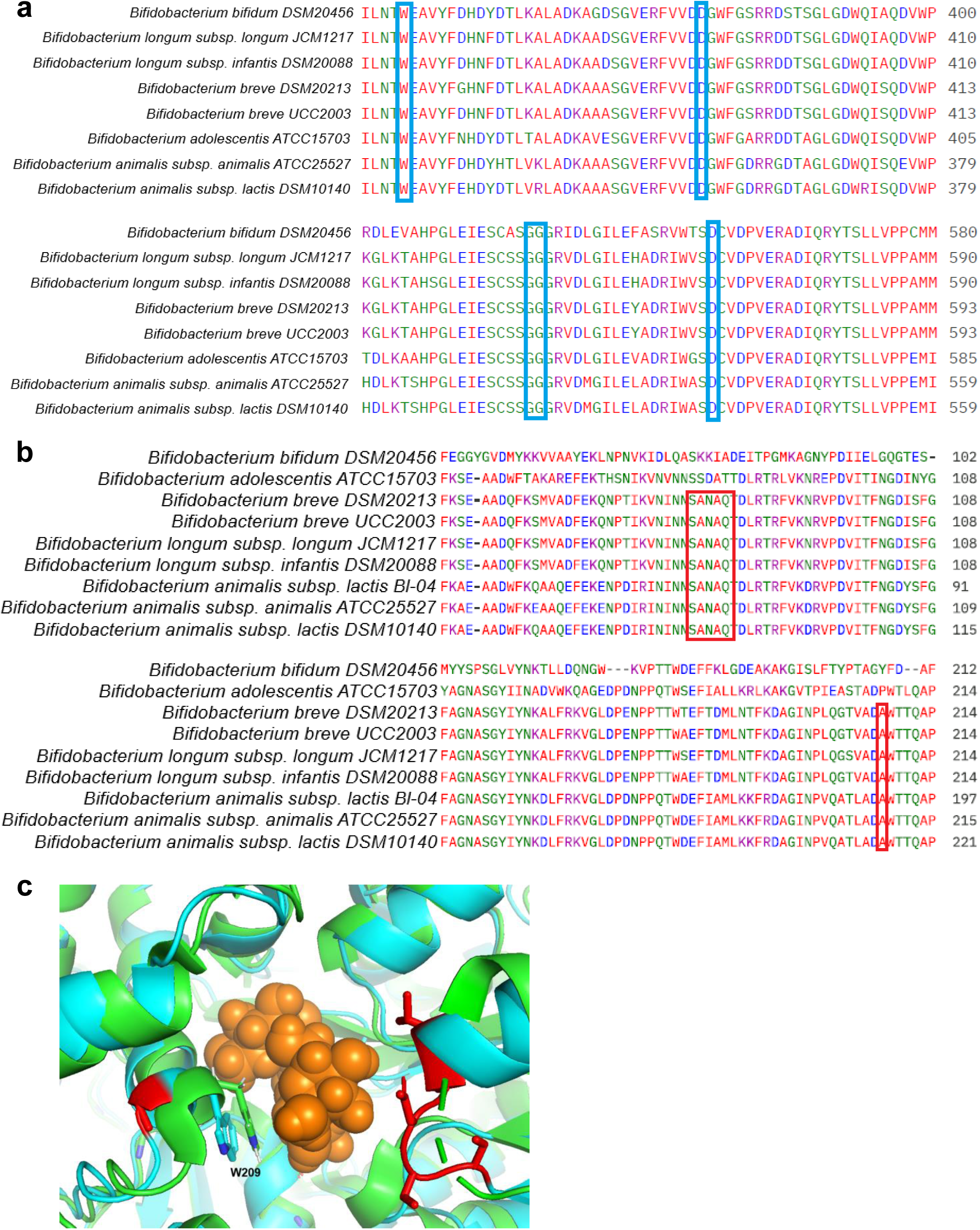
Raffinose utilization of *Bifidobacterium* is correlated with genetic capacity. **a**,**b**, Select regions of protein sequence alignments for putative RafA (**a**) and RafB (**b**) homologs are shown. **a,** Amino acids involved in direct binding to the galactose ring in PDB 2XN2, the closest homolog of *Bb* RafA available in the PDB, are highlighted in blue; all of these key raffinose-binding residues are strictly conserved in all bifidobacterial RafA sequences examined. **b,** Select amino acid residues that differ in *B. adolescentis* and *B. bifidum*, compared to the consensus RafB protein sequence from *Bifidobacterium* species that are rapidly rescued by raffinose *in vivo*, are boxed in red. **c**, An AlphaFold model of RafB from *B. adolescentis* (cyan) was aligned to PDB 4ZS9 (green), an experimentally-determined crystal structure of RafB from *B. lactis* Bl-04 in complex with raffinose^40^. Raffinose is shown as orange spheres. P208 in *B. adolescentis* RafB (red, left side) is predicted by the AlphaFold model to distort a local alpha-helical structure, resulting in a shift of residue W209 away from raffinose. Additionally, a short loop in *B. adolescentis* RafB, 79-SSDAT-83 (red, right side), diverges from the consensus sequence of *Bifidobacterium* species that are rapidly rescued by raffinose *in vivo*. The structural alignment shows that this loop is positioned at the entry of the raffinose binding pocket, consistent with a potential role in raffinose movement into or out of the binding pocket.

**Extended Data Fig. 5.**
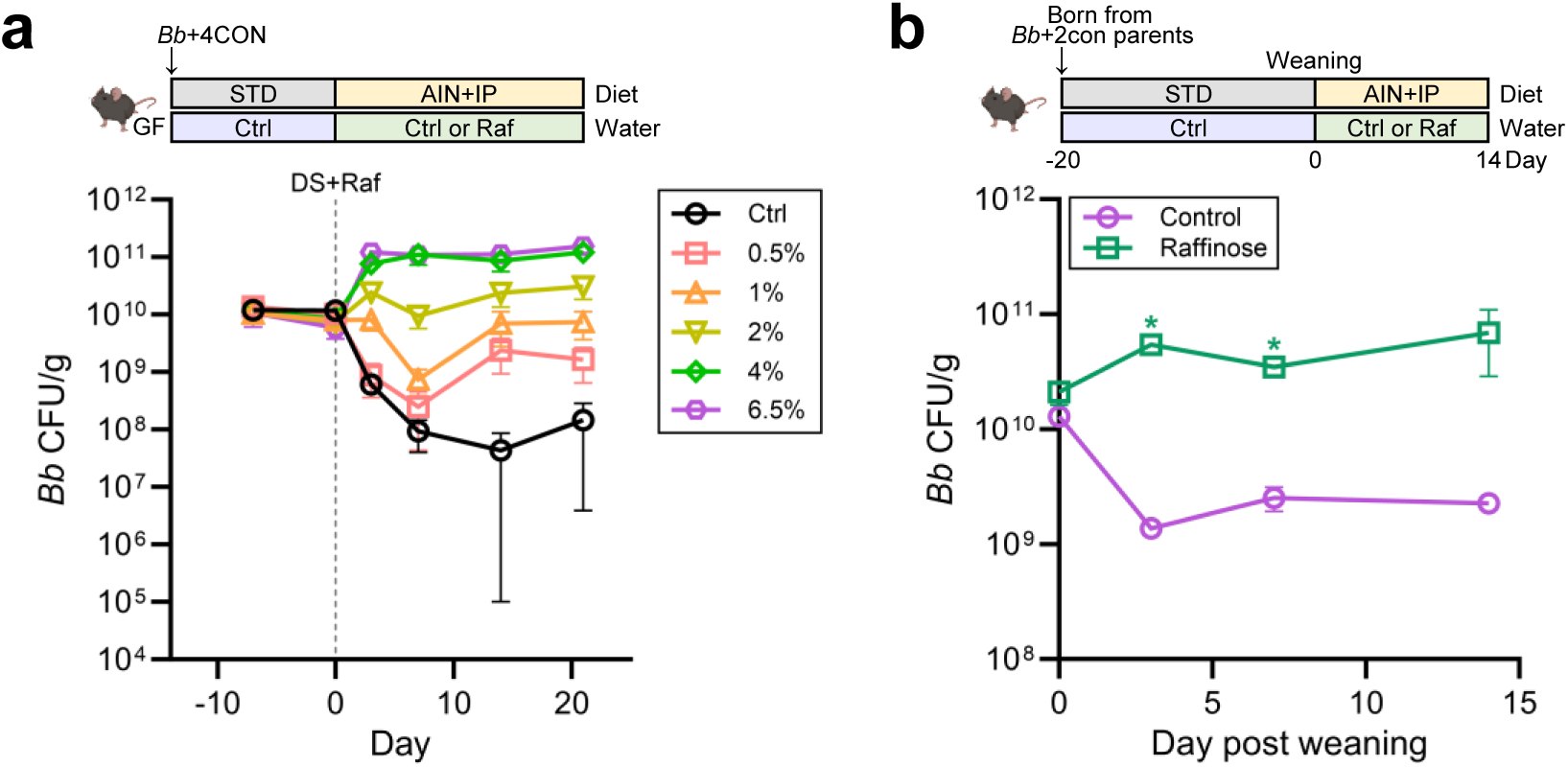
Raffinose rescues diet-induced loss of *Bb* colonization in a concentration-dependent but age-independent manner. **a**, Germ-free mice were colonized with *Bb*+4CON and maintained on STD for 2 weeks. Then, mice were switched to AIN+IP with or without raffinose-supplemented water at the indicated concentrations (% w/v). Fecal *Bb* CFUs were quantified over time by plating. Data represent *n* = 4-5 per group. **b**, Pups born from *Bb*+2CON-colonized parents were weaned at 20 days after birth. Both parents and pups were maintained on STD before weaning, and the pups were switched to AIN+IP with or without 4% (w/v) raffinose-supplemented water at weaning. Fecal *Bb* CFUs were quantified over time by plating. Data represent *n* = 2 per group. All data are shown as mean ± SEM and were compared by Student’s *t*-test (\**p* < 0.05).

**Extended Data Fig. 6.**
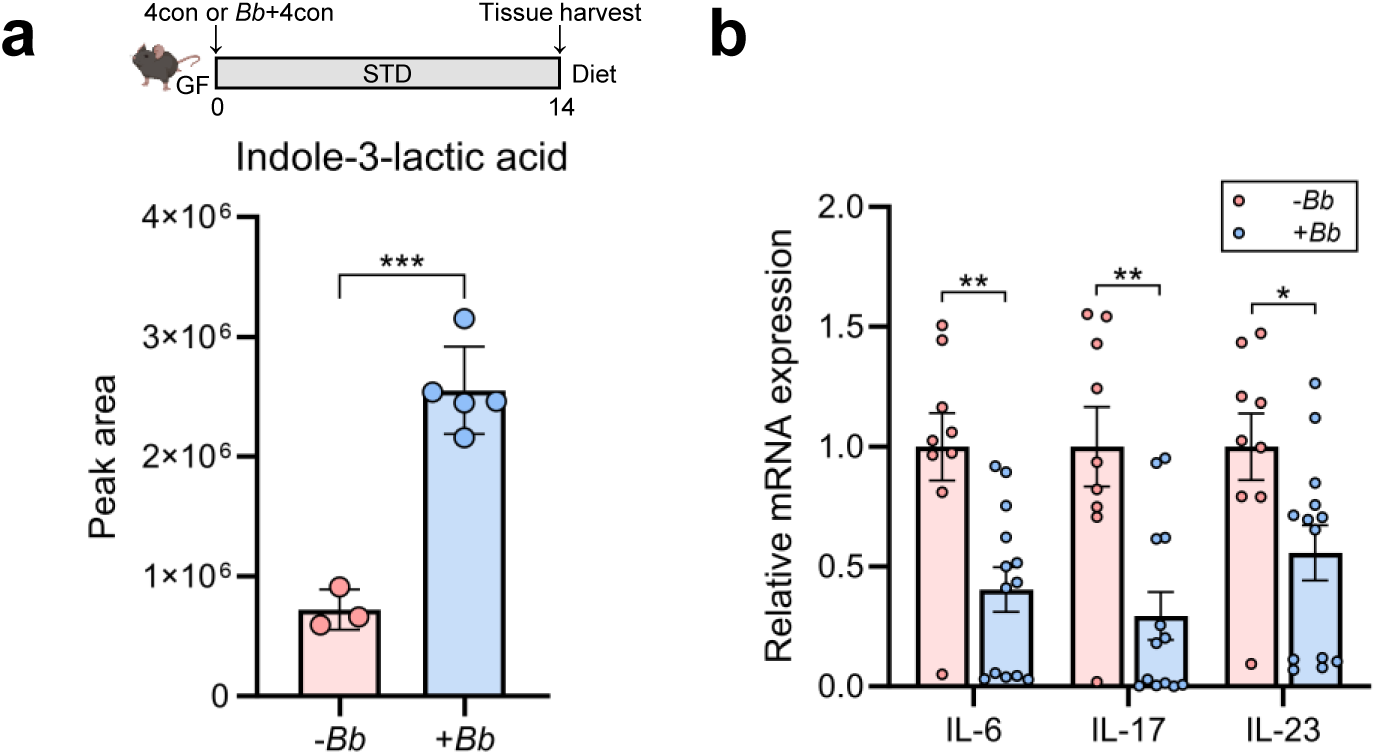
*Bb* exhibits anti-inflammatory effects in healthy consortium-colonized mice. Germ-free mice were colonized either with 4CON only or *Bb*+4CON and maintained on STD for 2 weeks. Then, mice were sacrificed and used to analyze host readouts. **a**, Cecal ILA levels were measured by metabolomics. Data represent *n* = 3 for -*Bb* and *n* = 5 for +*Bb*. **b**, Ileal expression of inflammatory cytokines was determined by qPCR and normalized to the control group. Data were pooled from three independent experiments (*n* = 9 for -*Bb*; *n* = 13 for +*Bb*). All data are shown as mean ± SEM and were compared by Student’s *t*-test (\**p* < 0.05; \*\**p* < 0.005; \*\*\**p* < 0.0005).

**Extended Data Fig. 7.**
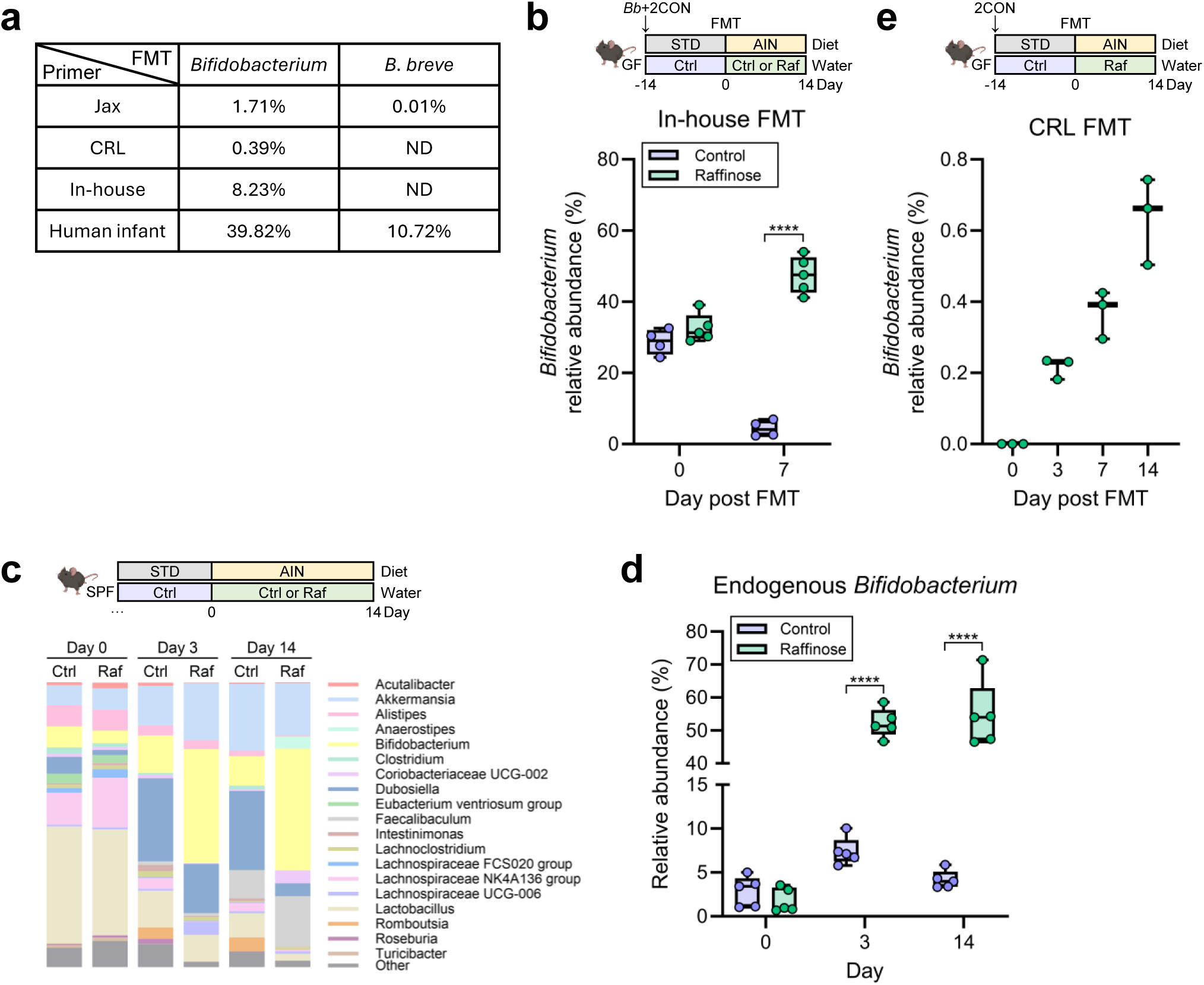
Raffinose supports intestinal persistence of pre-colonized *Bb* and endogenous *Bifidobacterium* in conventionalized and naïve SPF mice. **a**, The relative abundance of endogenous *Bifidobacterium* in fecal microbiota transfer (FMT) stocks derived from Jax, CRL, in-house SPF mice, or a human infant was analyzed by qPCR and normalized to total bacterial 16S rRNA levels. ND, not detected. **b**, Germ-free mice were colonized with *Bb*+2CON and maintained on STD for one week. Then, fecal microbiota from in-house SPF mice were introduced to induce microbiota diversification. Simultaneously, mice were switched to AIN with or without 6.5% (w/v) raffinose-supplemented water. The relative abundance of *Bifidobacterium* after FMT and diet switch was analyzed by qPCR and normalized to total bacterial 16S rRNA levels. Data represent *n* = 4 for control and *n* = 5 for raffinose. **c**,**d**, Naïve Jax SPF mice were maintained on STD for one week and then switched to AIN with or without 6.5% (w/v) raffinose-supplemented water. Fecal microbiota at day 0, 3, and 14 after diet switch were analyzed by 16S rRNA gene sequencing and differential abundance analysis was performed. Data represent *n* = 5 per group. **d**, Relative abundance of endogenous *Bifidobacterium* following diet switch. **e**, Germ-free mice were colonized with 2CON and maintained on STD for 2 weeks. Then, fecal microbiota from CRL SPF mice were introduced to induce microbiota diversification. Simultaneously, mice were switched to AIN with 6.5% (w/v) raffinose-supplemented water. The relative abundance of endogenous *Bifidobacterium* after FMT and diet switch was analyzed by qPCR and normalized to total bacterial 16S rRNA levels (*n* = 3). Data in **b**,**d**,**e** are shown as box-and-whisker plots with individual data points representing each mouse and were compared by Student’s *t*-test (\*\*\*\**p* < 0.0001).

**Extended Data Fig. 8.**
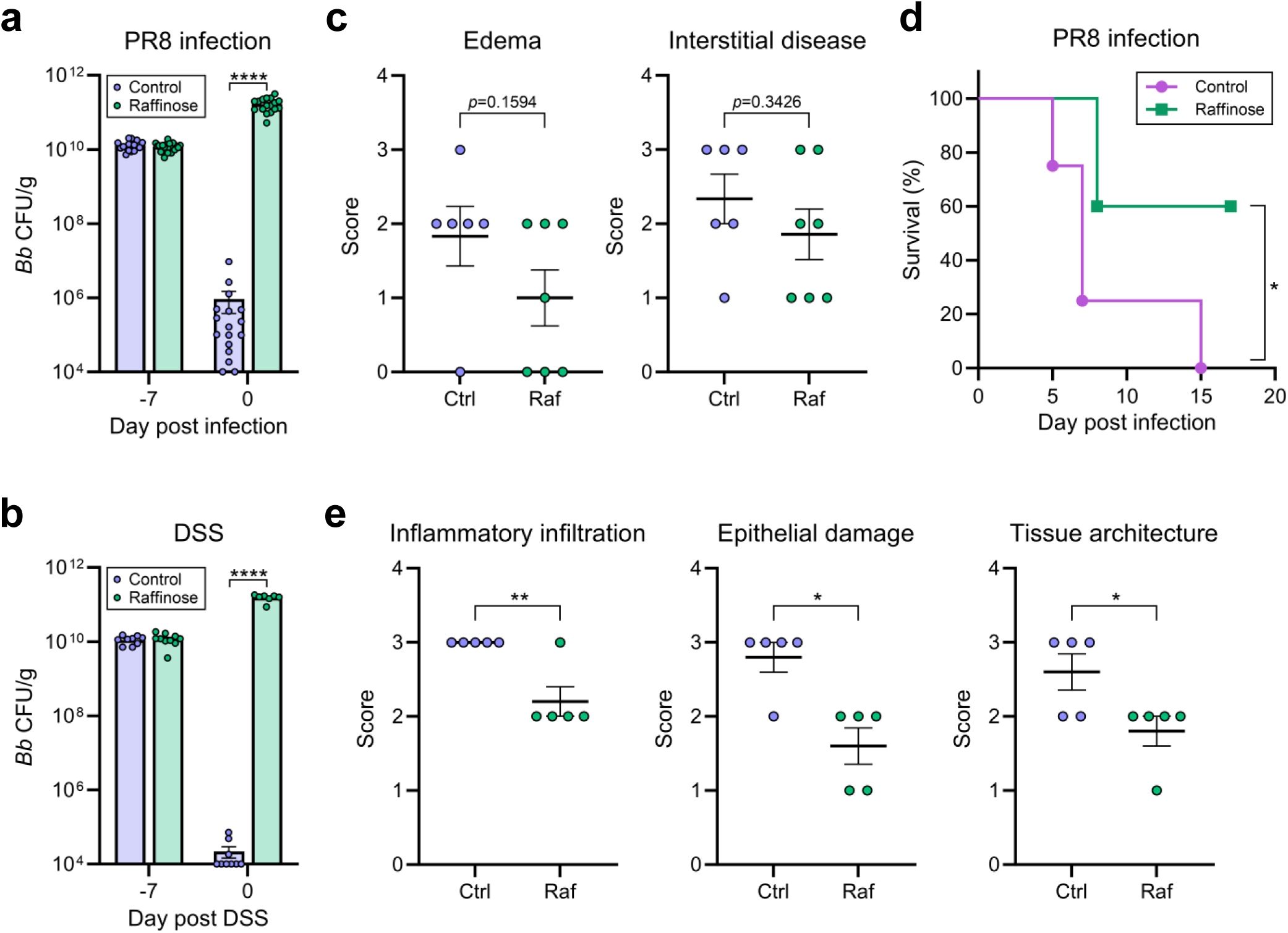
Raffinose improves disease outcomes in the influenza infection and DSS-induced colitis models in conventionalized mice. Germ-free mice were colonized with *Bb*+2CON and maintained on STD for one week. Then, mice received Jax FMT and were switched to AIN with or without 6.5% (w/v) raffinose-supplemented water. After an additional week, at day 0 mice were either infected intranasally with PR8 (25 PFU; **a**,**c**,**d**) or administered 2.5% (w/v) DSS for 4 days (**b**,**e**). **a**,**b**, Fecal *Bb* CFUs at the time of FMT and PR8 infection (**a**) or DSS administration (**b**) were quantified by plating. Data in **a** were pooled from four independent experiments (*n* = 17 for control; *n* = 19 for raffinose). Data in **b** were pooled from two independent experiments (*n* = 9 for control; *n* = 10 for raffinose). **c**, Pulmonary edema in airspaces and interstitial disease were evaluated in lung sections collected at day 6 post PR8 infection and quantified using an ordinal scoring system as described in Methods. Data represent *n* = 6 for control and *n* = 7 for raffinose from one independent experiment. **d**, Mouse survival following PR8 infection was monitored for 17 days (*n* = 4 for control; *n* = 5 for raffinose). Survival curves were compared using the log-rank (Mantel–Cox) test (\**p* < 0.05). **e**, Distal colon histology was evaluated in sections collected at day 5 post DSS administration. Inflammatory infiltration, epithelial damage, and tissue architecture disruption were scored separately as described in Methods. Data represent *n* = 5 per group from one independent experiment. Data in **a**-**c**,**e** are shown as mean ± SEM and were compared by Student’s *t*-test (\**p* < 0.05; \*\**p* < 0.005; \*\*\*\**p* < 0.0001).

**Extended Data Fig. 9.**
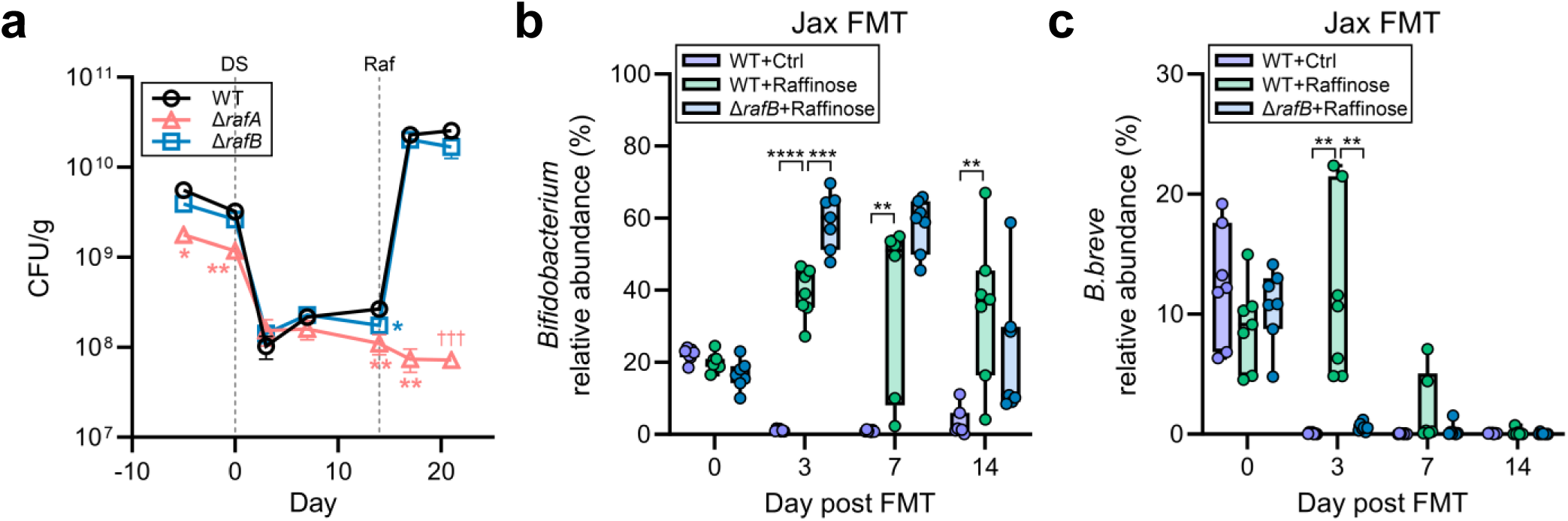
RafB contributes to the persistence of *B. breve* after microbiota diversification. **a**, Germ-free mice were mono-colonized with either one of *B. breve* UCC2003 WT, Δ*rafA*, or Δ*rafB* strain and maintained on STD for 10 days before being switched to AIN+IP. After 2 weeks on AIN+IP, mice were given drinking water supplemented with 6.5% (w/v) raffinose. Fecal CFUs of each strain were quantified over time by plating. Data were pooled from two independent experiments for WT (*n* = 7) and Δ*rafB* (*n* = 8) and from one independent experiment for Δ*rafA* (*n* = 3 for all time points except day 21, where *n* = 1). Mouse deaths are indicated with daggers. Data are shown as mean ± SEM and each group was compared to the WT group by Student’s *t*-test (\**p* < 0.05; \*\**p* < 0.005). **b**,**c**, Germ-free mice were colonized either with *B. breve* UCC2003 WT+2CON or Δ*rafB*+2CON and maintained on STD for 2 weeks. Then, fecal microbiota from Jax SPF mice was introduced by oral gavage to induce microbiota diversification. Simultaneously, mice were switched to AIN with or without 6.5% (w/v) raffinose-supplemented water. The relative abundance of total *Bifidobacterium*, including both endogenous mouse *Bifidobacterium* and the gavage *B. breve* UCC2003 strain (using a genus-specific primer set; **b**) and *B. breve* UCC2003 (using a species-specific primer set; **c**) after FMT and diet switch was analyzed by qPCR and normalized to total bacterial 16S rRNA levels. Data are shown as box-and-whisker plots with individual data points representing each mouse. Data were pooled from two independent experiments (*n* = 7 for WT+Ctrl, *n* = 6-7 for WT+Raf, *n* = 7 for Δ*rafB*+Raf) and compared by Student’s *t*-test (\*\**p* < 0.005; \*\*\**p* < 0.0005; \*\*\*\**p* < 0.0001).

## Reference

1 Stewart, C. J. et al. Temporal development of the gut microbiome in early childhood from the TEDDY study. Nature 562, 583–588 (2018). 10.1038/s41586-018-0617-x

2 Vatanen, T. et al. The human gut microbiome in early-onset type 1 diabetes from the TEDDY study. Nature 562, 589–594 (2018). 10.1038/s41586-018-0620-2

3 Charbonneau, M. R. et al. Sialylated Milk Oligosaccharides Promote Microbiota-Dependent Growth in Models of Infant Undernutrition. Cell 164, 859–871 (2016). 10.1016/j.cell.2016.01.024

4 Barratt, M. J. et al. Bifidobacterium infantis treatment promotes weight gain in Bangladeshi infants with severe acute malnutrition. Sci Transl Med 14, eabk1107 (2022). 10.1126/scitranslmed.abk1107

5 Sarkar, A. & Mandal, S. Bifidobacteria-Insight into clinical outcomes and mechanisms of its probiotic action. Microbiol Res 192, 159–171 (2016). 10.1016/j.micres.2016.07.001

6 Bessman, N. J. et al. Dendritic cell-derived hepcidin sequesters iron from the microbiota to promote mucosal healing. Science 368, 186–189 (2020). 10.1126/science.aau6481

7 Fukuda, S. et al. Bifidobacteria can protect from enteropathogenic infection through production of acetate. Nature 469, 543–547 (2011). 10.1038/nature09646

8 Odenwald, M. A. et al. Bifidobacteria metabolize lactulose to optimize gut metabolites and prevent systemic infection in patients with liver disease. Nat Microbiol 8, 2033–2049 (2023). 10.1038/s41564-023-01493-w

9 Sivan, A. et al. Commensal Bifidobacterium promotes antitumor immunity and facilitates anti-PD-L1 efficacy. Science 350, 1084–1089 (2015). 10.1126/science.aac4255

10 Mager, L. F. et al. Microbiome-derived inosine modulates response to checkpoint inhibitor immunotherapy. Science 369, 1481–1489 (2020). 10.1126/science.abc3421

11 Henrick, B. M. et al. Bifidobacteria-mediated immune system imprinting early in life. Cell 184, 3884–3898 e3811 (2021). 10.1016/j.cell.2021.05.030

12 Ryan, F. J. et al. Bifidobacteria support optimal infant vaccine responses. Nature 641, 456–464 (2025). 10.1038/s41586-025-08796-4

13 Stevens, J. et al. Microbiota-derived inosine programs protective CD8(+) T cell responses against influenza in newborns. Cell 188, 4239–4256 e4219 (2025). 10.1016/j.cell.2025.05.013

14 Arboleya, S., Watkins, C., Stanton, C. & Ross, R. P. Gut Bifidobacteria Populations in Human Health and Aging. Front Microbiol 7, 1204 (2016). 10.3389/fmicb.2016.01204

15 Derrien, M., Turroni, F., Ventura, M. & van Sinderen, D. Insights into endogenous Bifidobacterium species in the human gut microbiota during adulthood. Trends Microbiol 30, 940–947 (2022). 10.1016/j.tim.2022.04.004

16 Abdill, R. J. et al. Integration of 168,000 samples reveals global patterns of the human gut microbiome. Cell 188, 1100–1118 e1117 (2025). 10.1016/j.cell.2024.12.017

17 Sela, D. A. & Mills, D. A. Nursing our microbiota: molecular linkages between bifidobacteria and milk oligosaccharides. Trends Microbiol 18, 298–307 (2010). 10.1016/j.tim.2010.03.008

18 Marcobal, A. et al. Bacteroides in the infant gut consume milk oligosaccharides via mucus-utilization pathways. Cell Host Microbe 10, 507–514 (2011). 10.1016/j.chom.2011.10.007

19 Button, J. E. et al. Dosing a synbiotic of human milk oligosaccharides and B. infantis leads to reversible engraftment in healthy adult microbiomes without antibiotics. Cell Host Microbe 30, 712–725 e717 (2022). 10.1016/j.chom.2022.04.001

20 Carter, M. M. et al. A human milk oligosaccharide alters the microbiome, circulating hormones, and metabolites in a randomized controlled trial of older adults. Cell Rep Med 6, 102256 (2025). 10.1016/j.xcrm.2025.102256

21 Li, F. et al. Cardiometabolic benefits of a non-industrialized-type diet are linked to gut microbiome modulation. Cell 188, 1226–1247 e1218 (2025). 10.1016/j.cell.2024.12.034

22 Dinoto, A. et al. Population dynamics of Bifidobacterium species in human feces during raffinose administration monitored by fluorescence in situ hybridization-flow cytometry. Appl Environ Microbiol 72, 7739–7747 (2006). 10.1128/AEM.01777-06

23 Kleessen, B. et al. Jerusalem artichoke and chicory inulin in bakery products affect faecal microbiota of healthy volunteers. Br J Nutr 98, 540–549 (2007). 10.1017/S0007114507730751

24 Vandeputte, D. et al. Prebiotic inulin-type fructans induce specific changes in the human gut microbiota. Gut 66, 1968–1974 (2017). 10.1136/gutjnl-2016-313271

25 Nicolucci, A. C. et al. Prebiotics Reduce Body Fat and Alter Intestinal Microbiota in Children Who Are Overweight or With Obesity. Gastroenterology 153, 711–722 (2017). 10.1053/j.gastro.2017.05.055

26 Magne, F. et al. Effects on faecal microbiota of dietary and acidic oligosaccharides in children during partial formula feeding. J Pediatr Gastroenterol Nutr 46, 580–588 (2008). 10.1097/MPG.0b013e318164d920

27 Jaeggi, T. et al. Iron fortification adversely affects the gut microbiome, increases pathogen abundance and induces intestinal inflammation in Kenyan infants. Gut 64, 731–742 (2015). 10.1136/gutjnl-2014-307720

28 Laursen, M. F. & Roager, H. M. Human milk oligosaccharides modify the strength of priority effects in the Bifidobacterium community assembly during infancy. ISME J 17, 2452–2457 (2023). 10.1038/s41396-023-01525-7

29 Maldonado-Gomez, M. X. et al. Stable Engraftment of Bifidobacterium longum AH1206 in the Human Gut Depends on Individualized Features of the Resident Microbiome. Cell Host Microbe 20, 515–526 (2016). 10.1016/j.chom.2016.09.001

30 Grimm, V., Radulovic, K. & Riedel, C. U. Colonization of C57BL/6 Mice by a Potential Probiotic Bifidobacterium bifidum Strain under Germ-Free and Specific Pathogen-Free Conditions and during Experimental Colitis. PLoS One 10, e0139935 (2015). 10.1371/journal.pone.0139935

31 Heiss, B. E. et al. Bifidobacterium catabolism of human milk oligosaccharides overrides endogenous competitive exclusion driving colonization and protection. Gut Microbes 13, 1986666 (2021). 10.1080/19490976.2021.1986666

32 Lubin, J. B. et al. Arresting microbiome development limits immune system maturation and resistance to infection in mice. Cell Host Microbe 31, 554–570 e557 (2023). 10.1016/j.chom.2023.03.006

33 Shao, Y. et al. Primary succession of Bifidobacteria drives pathogen resistance in neonatal microbiota assembly. Nat Microbiol 9, 2570–2582 (2024). 10.1038/s41564-024-01804-9

34 Sun, S. et al. Bifidobacterium alters the gut microbiota and modulates the functional metabolism of T regulatory cells in the context of immune checkpoint blockade. Proc Natl Acad Sci U S A 117, 27509–27515 (2020). 10.1073/pnas.1921223117

35 Reeves, P. G., Rossow, K. L. & Lindlauf, J. Development and testing of the AIN-93 purified diets for rodents: results on growth, kidney calcification and bone mineralization in rats and mice. J Nutr 123, 1923–1931 (1993). 10.1093/jn/123.11.1923

36 Zomer, A. et al. An interactive regulatory network controls stress response in Bifidobacterium breve UCC2003. J Bacteriol 191, 7039–7049 (2009). 10.1128/JB.00897-09

37 Zuo, F. et al. Transcriptomic analysis of Bifidobacterium longum subsp. longum BBMN68 in response to oxidative shock. Sci Rep 8, 17085 (2018). 10.1038/s41598-018-35286-7

38 Kennedy, M. S. et al. Dynamic genetic adaptation of Bacteroides thetaiotaomicron during murine gut colonization. Cell Rep 42, 113009 (2023). 10.1016/j.celrep.2023.113009

39 Shiver, A. L. et al. Genome-scale resources in the infant gut symbiont Bifidobacterium breve reveal genetic determinants of colonization and host-microbe interactions. Cell 188, 2003–2021 e2019 (2025). 10.1016/j.cell.2025.02.010

40 Ejby, M. et al. An ATP Binding Cassette Transporter Mediates the Uptake of alpha-(1,6)-Linked Dietary Oligosaccharides in Bifidobacterium and Correlates with Competitive Growth on These Substrates. J Biol Chem 291, 20220–20231 (2016). 10.1074/jbc.M116.746529

41 Brugiroux, S. et al. Genome-guided design of a defined mouse microbiota that confers colonization resistance against Salmonella enterica serovar Typhimurium. Nat Microbiol 2, 16215 (2016). 10.1038/nmicrobiol.2016.215

42 Cheng, A. G. et al. Design, construction, and in vivo augmentation of a complex gut microbiome. Cell 185, 3617–3636 e3619 (2022). 10.1016/j.cell.2022.08.003

43 Zhang, Q. et al. Lactobacillus plantarum-derived indole-3-lactic acid ameliorates colorectal tumorigenesis via epigenetic regulation of CD8(+) T cell immunity. Cell Metab 35, 943–960 e949 (2023). 10.1016/j.cmet.2023.04.015

44 Laursen, M. F. et al. Bifidobacterium species associated with breastfeeding produce aromatic lactic acids in the infant gut. Nat Microbiol 6, 1367–1382 (2021). 10.1038/s41564-021-00970-4

45 Zhou, L. et al. Group 3 innate lymphoid cells produce the growth factor HB-EGF to protect the intestine from TNF-mediated inflammation. Nat Immunol 23, 251–261 (2022). 10.1038/s41590-021-01110-0

46 Zheng, B. et al. Bifidobacterium breve attenuates murine dextran sodium sulfate-induced colitis and increases regulatory T cell responses. PLoS One 9, e95441 (2014). 10.1371/journal.pone.0095441

47 Shao, Y. et al. Genomic atlas of Bifidobacterium infantis and B. longum informs infant probiotic design. Cell 189, 1854–1873 e1817 (2026). 10.1016/j.cell.2026.01.007

48 Gabryszewski, S. J. et al. Guidelines for Early Food Introduction and Patterns of Food Allergy. Pediatrics 156 (2025). 10.1542/peds.2024-070516

49 Du Toit, G. et al. Randomized trial of peanut consumption in infants at risk for peanut allergy. N Engl J Med 372, 803–813 (2015). 10.1056/NEJMoa1414850

50 Yamashita, H. et al. Impact of orally-administered oligosaccharides in a murine model of food allergy. Journal of Functional Foods 85 (2021/10/01). 10.1016/j.jff.2021.104643

51 Dobin, A. et al. STAR: ultrafast universal RNA-seq aligner. Bioinformatics 29, 15–21 (2013). 10.1093/bioinformatics/bts635

52 Anders, S., Pyl, P. T. & Huber, W. HTSeq--a Python framework to work with high-throughput sequencing data. Bioinformatics 31, 166–169 (2015). 10.1093/bioinformatics/btu638

53 Love, M. I., Huber, W. & Anders, S. Moderated estimation of fold change and dispersion for RNA-seq data with DESeq2. Genome Biol 15, 550 (2014). 10.1186/s13059-014-0550-8

54 O’Connell Motherway, M., et al. Functional genome analysis of Bifidobacterium breve UCC2003 reveals type IVb tight adherence (Tad) pili as an essential and conserved host-colonization factor. Proc Natl Acad Sci U S A 108, 11217–11222 (2011). 10.1073/pnas.1105380108

55 Lu, P., Shih, C. & Qi, H. Ephrin B1-mediated repulsion and signaling control germinal center T cell territoriality and function. Science 356 (2017). 10.1126/science.aai9264

56 Altschul, S. F. et al. Gapped BLAST and PSI-BLAST: a new generation of protein database search programs. Nucleic Acids Res 25, 3389–3402 (1997). 10.1093/nar/25.17.3389

57 Sievers, F. et al. Fast, scalable generation of high-quality protein multiple sequence alignments using Clustal Omega. Mol Syst Biol 7, 539 (2011). 10.1038/msb.2011.75

58 Varadi, M. et al. AlphaFold Protein Structure Database: massively expanding the structural coverage of protein-sequence space with high-accuracy models. Nucleic Acids Res 50, D439–D444 (2022). 10.1093/nar/gkab1061

59 Martin, M. Cutadapt removes adapter sequences from high-throughput sequencing reads. EMBnet.journal 17 (2011/05/02). 10.14806/ej.17.1.200

60 Callahan, B. J. et al. DADA2: High-resolution sample inference from Illumina amplicon data. Nat Methods 13, 581–583 (2016). 10.1038/nmeth.3869

61 Quast, C. et al. The SILVA ribosomal RNA gene database project: improved data processing and web-based tools. Nucleic Acids Res 41, D590–596 (2013). 10.1093/nar/gks1219

62 Bokulich, N. A. et al. Optimizing taxonomic classification of marker-gene amplicon sequences with QIIME 2’s q2-feature-classifier plugin. Microbiome 6, 90 (2018). 10.1186/s40168-018-0470-z

63 Helf, M. J., Fox, B. W., Artyukhin, A. B., Zhang, Y. K. & Schroeder, F. C. Comparative metabolomics with Metaboseek reveals functions of a conserved fat metabolism pathway in C. elegans. Nat Commun 13, 782 (2022). 10.1038/s41467-022-28391-9

64 Tautenhahn, R., Bottcher, C. & Neumann, S. Highly sensitive feature detection for high resolution LC/MS. BMC Bioinformatics 9, 504 (2008). 10.1186/1471-2105-9-504

65 Sun, H. et al. Autologous fecal microbiota transplantation restores the infant gut microbiome and metabolome after antibiotics: a case report. mBio, e0071126 (2026). 10.1128/mbio.00711-26

66 Meyerholz, D. K. & Beck, A. P. Principles and approaches for reproducible scoring of tissue stains in research. Lab Invest 98, 844–855 (2018). 10.1038/s41374-018-0057-0

67 Aeffner, F. et al. Scientific and Regulatory Policy Committee Points to Consider*: Sample Selection, Assay Design, Data Generation and Interpretation, and Reporting Practices for Chromogenic Immunohistochemical (IHC) Assays in Nonclinical Drug Development. Toxicol Pathol, 1926233251393154 (2025). 10.1177/01926233251393154

68 Wong, L. R. et al. Eicosanoid signalling blockade protects middle-aged mice from severe COVID-19. Nature 605, 146–151 (2022). 10.1038/s41586-022-04630-3

69 Remke, M. et al. Histomorphological scoring of murine colitis models: A practical guide for the evaluation of colitis and colitis-associated cancer. Exp Mol Pathol 140, 104938 (2024). 10.1016/j.yexmp.2024.104938

